# Transcriptional profiling reveals extraordinary diversity among skeletal muscle tissues

**DOI:** 10.1101/216317

**Authors:** Erin E Terry, Xiping Zhang, Christy Hoffmann, Laura D Hughes, Scott A Lewis, Jiajia Li, Lance Riley, Nicholas F Lahens, Ming Gong, Francisco Andrade, Karyn A Esser, Michael E. Hughes

## Abstract

Skeletal muscle comprises a family of diverse tissues with highly specialized morphology, function, and metabolism. Many acquired diseases – including HIV, COPD, cancer cachexia, critical illness myopathy, and sepsis – affect specific muscles while sparing others. Even monogenic muscular dystrophies tend to selectively affect certain muscle groups, despite their causative genetic mutations being present in all tissues. These observations suggest that factors intrinsic to muscle tissues influence their susceptibility to various disease mechanisms. Nevertheless, most studies have not addressed transcriptional diversity among skeletal muscles. Here we use RNA sequencing (RNA-seq) to profile global mRNA expression in a wide array of skeletal, smooth, and cardiac muscle tissues from mice and rats. Our data set, MuscleDB, reveals extensive transcriptional diversity, with greater than 50% of transcripts differentially expressed among skeletal muscle tissues. This diversity is only partly explained by fiber type composition and developmental history, suggesting that specialized transcriptional profiles establish the functional identity of muscle tissues. We find conservation in the transcriptional profiles across species as well as between males and females, indicating that these data may be useful in predicting gene expression in related species. Notably, thousands of differentially expressed genes in skeletal muscle are associated with human disease, and hundreds of these genes encode targets of drugs on the market today. We detect mRNA expression of hundreds of putative myokines that may underlie the endocrine functions of skeletal muscle. In addition to demonstrating the intrinsic diversity of skeletal muscles, these data provide a resource for generating testable hypotheses regarding the mechanisms that establish differential disease susceptibility in muscle.

**Significance Statement:** Skeletal muscles are a diverse family of tissues with a common contractile function but divergent morphology, development, and metabolism. One need only reflect on the different functions of limb muscles and the diaphragm to realize the highly specialized nature of these tissues. Nevertheless, every study of global gene expression has analyzed at most one representative skeletal muscle. Here we measure gene expression from 11 different skeletal muscles in mice and rats. We show that there is no such thing as a representative skeletal muscle, as gene expression profiles vary widely among the tissues analyzed. These data are an important resource for pharmacologists, tissue engineers, and investigators studying the mechanisms of cellular specialization.

## Introduction

Gene expression atlases have made enormous contributions to our understanding of genetic regulatory mechanisms. The field of functional genomics was set in motion by the completion of the Human Genome Project and the coincident development of high throughput gene expression profiling technologies. The overriding goals of this field are to understand how genes and proteins interact at a whole-genome scale and to define how these interactions change across time, space, and different disease states. The development of SymAtlas was an early, deeply influential effort to address these questions (1). Custom microarrays were used to systematically profile mRNA expression in dozens of tissues and cell lines from humans and mice. Besides describing tissue-specific expression patterns, these data provided essential insights into the relationship between chromosomal structure and transcriptional regulation (2).

Related approaches have had a similarly high impact on biomedical research. For example, microRNA expression throughout mammalian tissues was described in an expression atlas that has revolutionized the study of regulatory RNAs (3). Our lab has contributed to this growing literature with the creation of CircaDB, a database of tissue-specific mRNA rhythms in mice (4, 5). Taken together, these projects demonstrate that publically available functional genomics data have enduring value as a resource for the research community.

Nevertheless, previous gene expression atlases have largely ignored skeletal muscle. CircaDB includes gene expression data from the heart and whole calf muscle, but it does not distinguish between their constituent tissues. Similarly, SymAtlas profiles nearly 100 different tissues, but there is only a single representative sample for either cardiac or skeletal muscle. The microRNA Atlas includes over 250 human, mouse, and rat tissues; however, skeletal muscle is entirely absent from these data. More recent human gene expression atlases have similar biases (6–8). Many studies have compared muscle-specific gene expression in different tissues (9), fiber-types (10), or disease states (11, 12), but on the whole, there is no systematic analysis of transcriptional diversity in skeletal muscle.

This gap in the literature is problematic since skeletal muscle comprises a remarkably diverse group of tissues. Skeletal muscle groups originate from different developmental origins, and they have characteristic morphological specializations (13–15). Their physiological functions are similarly diversified. For example, extraocular muscles govern precise eye movements, the diaphragm drives rhythmic breathing, and limb skeletal muscles are involved in either fast bursts of motion or sustained contractions underlying posture.

These intrinsic differences contribute to differential susceptibility of muscle groups to injury and disease. For example, there are six major classes of muscular dystrophy, and each one afflicts a characteristic pattern of skeletal and cardiac muscle tissues (16, 17). Since the causative mutations underlying congenital muscular dystrophies are germline and present in all cells, this observation indicates that there are properties intrinsic to different muscle tissues that regulate their sensitivity or resistance to different pathological mechanisms.

Acquired diseases show similar specificity in which muscle tissues they affect and which they spare. Patients with chronic obstructive pulmonary disorder (COPD) typically have pronounced myopathy in their quadriceps and dorsiflexors (18–20). Critical illness myopathy, a debilitating condition caused by mechanical ventilation and steroid treatment, causes muscle weakness in limb and respiratory muscles while sparing facial muscles (21–23). Cancer cachexia (24), HIV (25), and sepsis (26) cause muscle wasting, typically affecting fast twitch fibers more severely than slow twitch fibers. In fact, histologically identical muscle fibers show widely divergent responses to injury and disease depending on the muscle group in which they reside (16, 27). Taken together, these observations strongly suggest that the intrinsic diversity of skeletal muscle has important consequences for human health and disease. However, the mechanisms through which disease susceptibility and functional specialization are established are unknown.

Here we present the first systematic examination of transcriptional programming in different skeletal muscle tissues. We find that more than 50% of transcripts are differentially expressed among skeletal muscle tissues, an observation that cannot be explained by fiber type composition or developmental history alone. We show conservation of gene expression profiles across species and sexes, suggesting that these data may reveal conserved functional elements relevant to human health. Finally, we discuss how this unique data set may be applied to the study of disease, particularly regarding muscular dystrophy and regenerative medicine.

## Results

To determine which skeletal muscle tissues are of the broadest general interest, we distributed a Google poll to leading investigators in the skeletal muscle field. Having recorded over 100 individual responses, we selected 11 mouse skeletal muscle tissues (**Table 1**) for study in order to span the functional, developmental, and anatomical diversity of skeletal muscles (**Figure S1**). To hedge against selection bias from the investigators asked to vote in the poll, we cross-correlated these results with papers indexed in NCBI’s PubMed (**Figure S1**). In addition to these 11 mouse skeletal muscle tissues, we also identified representative smooth and cardiac muscle tissues for collection from mouse, and two skeletal muscle tissues (*EDL* and *soleus*) from male and female rats to permit inter-species and inter-sex comparisons (**Table 1**).

**Table 1:** The biological samples collected in this study and the total number of aligned RNA-seq reads.

| **Tissue** | **Type** | **Species** | **Sex** | **Replicates** | **Aligned Reads**<br>**(Millions)** |
| --- | --- | --- | --- | --- | --- |
| Total Aorta | Smooth | Mouse | Male | 6 | 205.2 |
| Abdominal Aorta | Smooth | Mouse | Male | 6 | 219.7 |
| Thoracic Aorta | Smooth | Mouse | Male | 6 | 214.0 |
| Atria | Cardiac | Mouse | Male | 6 | 169.0 |
| Left Ventricle | Cardiac | Mouse | Male | 6 | 193.9 |
| Right Ventricle | Cardiac | Mouse | Male | 6 | 176.2 |
| Diaphragm | Skeletal | Mouse | Male | 6 | 159.1 |
| EDL | Skeletal | Mouse | Male | 6 | 176.8 |
| Extraocular | Skeletal | Mouse | Male | 6 | 171.7 |
| FDB | Skeletal | Mouse | Male | 6 | 202.5 |
| Masseter | Skeletal | Mouse | Male | 6 | 216.2 |
| Plantaris | Skeletal | Mouse | Male | 6 | 185.1 |
| Soleus | Skeletal | Mouse | Male | 6 | 172.2 |
| Tongue | Skeletal | Mouse | Male | 6 | 210.4 |
| Gastrocnemius | Skeletal | Mouse | Male | 6 | 276.1 |
| Quadriceps | Skeletal | Mouse | Male | 6 | 275.6 |
| Tibialis Anterior | Skeletal | Mouse | Male | 6 | 276.6 |
| EDL | Skeletal | Rat | Male | 6 | 226.4 |
| EDL | Skeletal | Rat | Female | 6 | 206.8 |
| Soleus | Skeletal | Rat | Male | 6 | 261.2 |
| Soleus | Skeletal | Rat | Female | 6 | 243.0 |
| Average |  |  |  |  | 211.3 |
| Total |  |  |  |  | 4,437.7 |

Adult mice and rats were sacrificed and whole muscle tissues were dissected to include the entire muscle body from tendon to tendon. Each sample included six biological replicates from three independent animals apiece. RNA was purified from these tissues, and RNA-seq was used to measure global gene expression (**Methods and Table S1**). On average, every muscle sample was covered by greater than 200 million aligned short nucleotide reads, for a grand total of over 4.4 billion aligned reads in the entire data set (**Table 1**). Empirical simulations indicate that this experimental design reaches saturation with respect to identification of expressed exons (**Figure S2**). Pairwise comparisons of each replicate sample further indicate a high degree of reproducibility in expression level measurements among biological replicates (median R^2^ value > 0.93, **Figure S3**).

Over 80% of transcripts encoded in the genome are expressed in at least one skeletal muscle (**Figure 1A**). Comparing the transcripts expressed in all tissues identifies a core group of ~21,000 transcripts found in every skeletal, cardiac, and smooth muscle (**Figure 1B**). Presumably, these mRNAs include the minimal set of genes required for a cell to generate contractile force. Differential expression analysis shows that even at extremely stringent (q < 10^-6^) statistical thresholds, 55.2% of mouse transcripts are differentially expressed among skeletal muscle tissues (**Figure 1C**). Phrased differently, over half of all transcripts are statistically different when comparing mRNA expression among the 11 mouse skeletal muscles in this study. To validate a subset of these data, we selected 10 genes differentially expressed between *EDL* and *soleus* across a range of different fold changes and performed quantitative PCR (qPCR) on independent biological samples. All 10 replicated, and manual examination of internal controls for cardiac and smooth muscle agreed with *a priori* expectations (**Figure S4**). To explore similarity between tissues, we calculated a pairwise Euclidean distance between every tissue (**Figure S5**). From these data, we generated a dendrogram that clusters tissues based on overall transcriptome similarity (**Figure 1D**). Smooth and cardiac tissues clustered together as expected, and skeletal muscles were found in two primary clusters, presumably based on the proportion of fast twitch fibers they include (i.e. *Myosin heavy chain 4* (*Myh4*) expression). We then calculated the proportion of differentially expressed transcripts in every pairwise tissue comparison (**Figure 1E**). Some surprising observations emerged from this analysis. For example, *masseter,* a head/neck muscle, is more similar to limb muscles like *EDL* than muscles that share a similar developmental history, such as the tongue. In contrast, the *flexor digitorm brevis* (*FDB*), a muscle necessary for flexing the toes, is more similar to extraocular muscles than limb muscles. On average, 13% of transcripts are differentially expressed between any two skeletal muscles, with a maximum of 36.5% and a minimum less than 1%.

**Figure 1:**
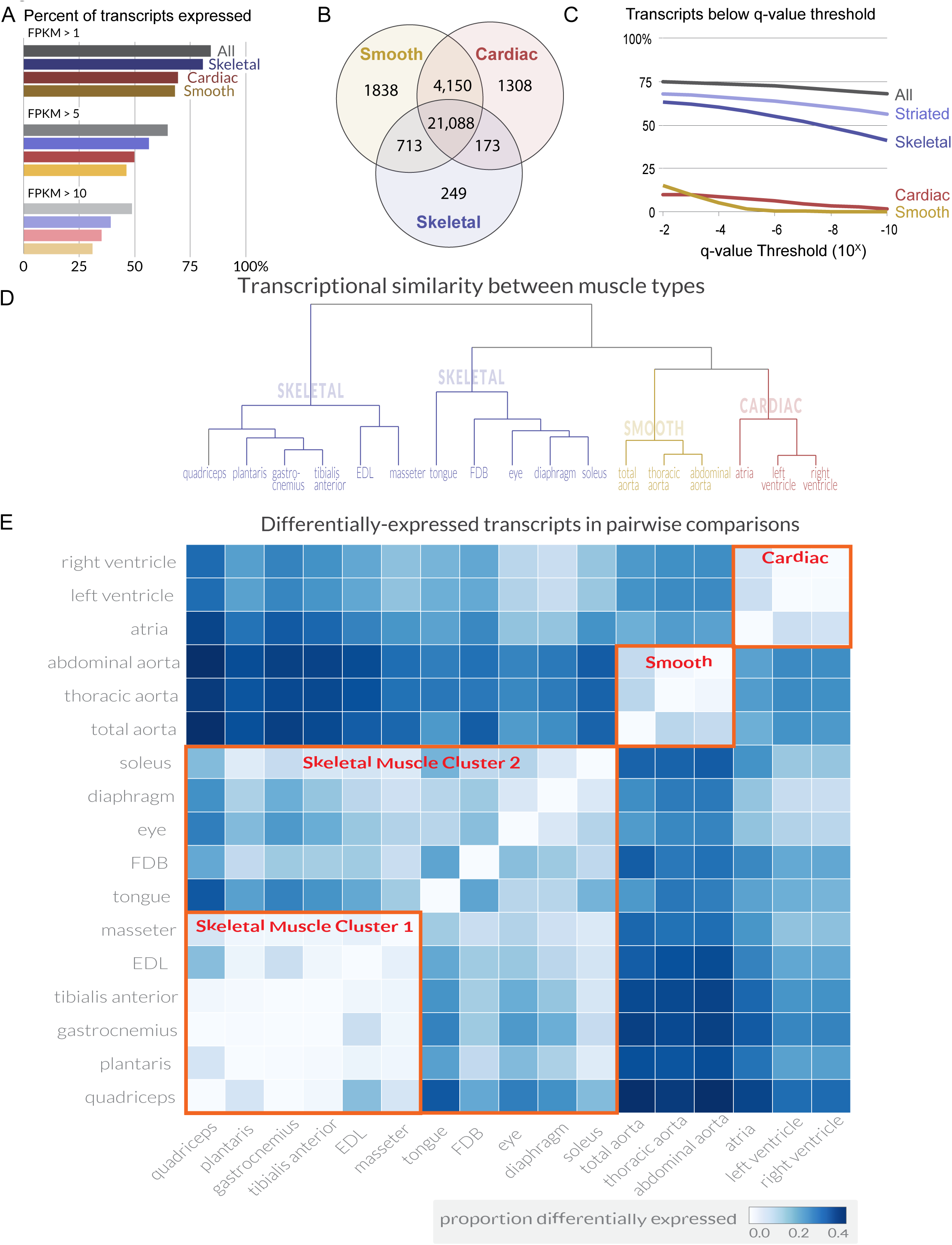
Transcriptome profiling reveals extensive gene expression differences among muscle tissues. (A) The percent of all transcripts detected as being expressed in different classes of tissues is shown as a bar graph. >80% of all transcripts are detectably expressed in at least one skeletal muscle tissue. (B) The number of transcripts expressed (FPKM > 1) in *every* cardiac, smooth, or skeletal muscle tissue is shown as a Venn diagram. A core of ~21,000 transcripts is expressed in every contractile tissue. (C) The percent of transcripts showing differential expression between tissues is shown at different false-discovery rate (q-value) thresholds. (D) The overall similarity of transcriptional profiles in different tissues is displayed as a dendrogram. Notably, the three major classes of muscle (smooth, cardiac, and skeletal) cluster together as expected. Striated muscle refers to skeletal plus cardiac muscles. (E) The number of differentially expressed transcripts (q < 0.01, fold change > 2) in pairwise comparisons is shown as a heat map. Red boxes indicate clustering by similarity of (1) cardiac muscle, (2) smooth muscle, and (3) two different clusters of skeletal muscle.

Skeletal muscle is comprised of myofibers which are typically classified into one of four types in mice based on their *Myh* expression (28). We used our data to quantify the relative abundance of different fiber types across skeletal muscle tissues (**Figure 2A**). These results were cross-correlated with legacy data from histological studies, showing close agreement (**Figure S6**). One hypothesis is that fiber type composition (i.e. the relative amounts of fast versus slow twitch fibers) establishes the observed diversity of gene expression profiles. To test this, we clustered skeletal muscle tissues based on similarity of *Myh* expression. The resulting dendrograms are grossly similar to those based on global gene expression (**Figure 2B**; compare with **Figure 1D**), as predominantly fast twitch muscles expressing high levels of *Myh4* (Type IIB fibers) tend to cluster together. Nevertheless, clustering within the two major skeletal muscle groups reveals important differences, such as the high similarity of diaphragm and *FDB* based on *Myh* expression, but their dissimilarity based on global gene expression (**Figure 1E**). Similarly, the clustering of tongue and extraocular eye muscles varies considerably depending on whether *Myh* or global gene expression establishes the pairwise Euclidean distance.

**Figure 2:**
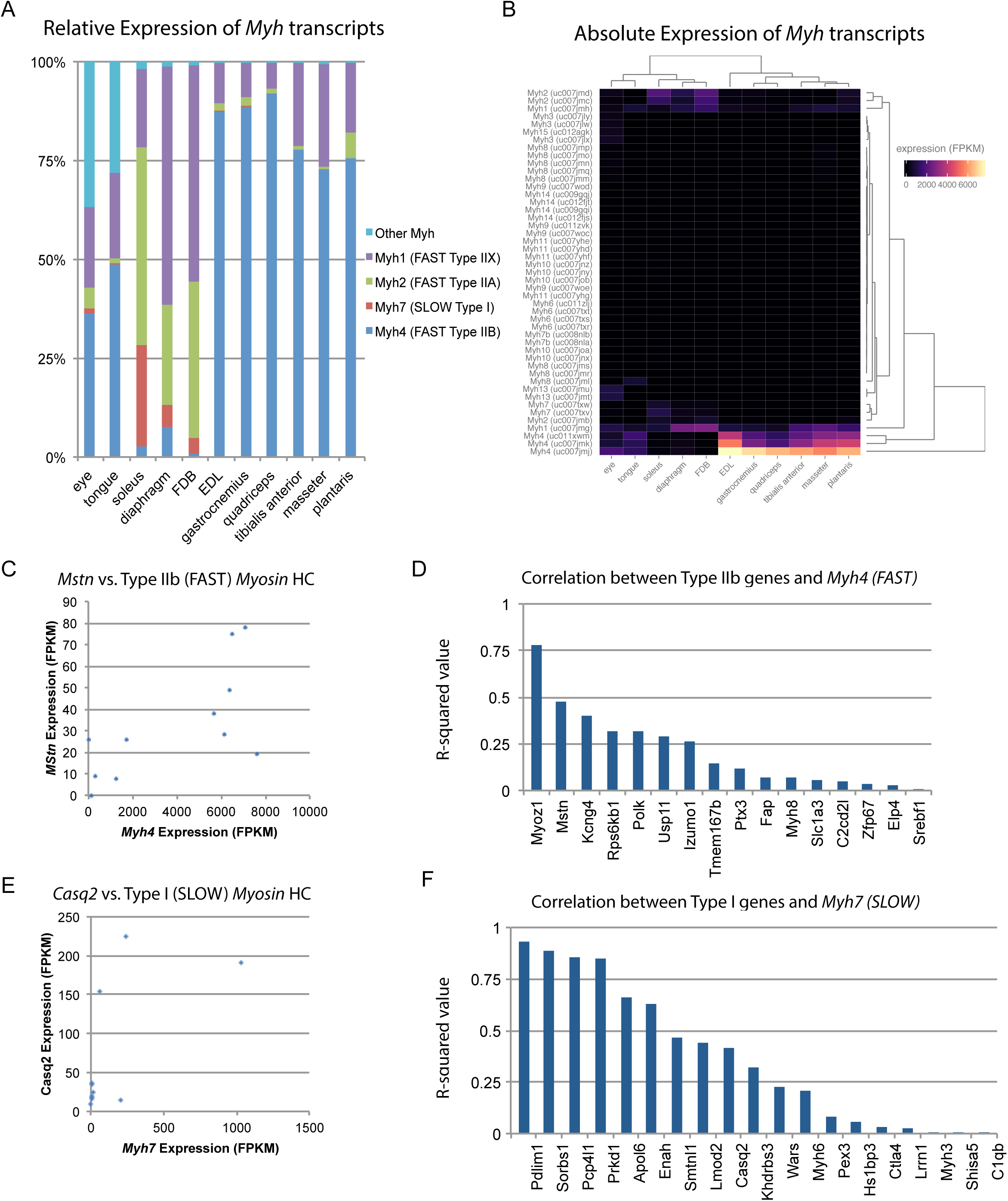
Fiber-type composition can explain some, but not all, transcriptional variability between skeletal muscle tissues. (A) Relative expression of *Myosin heavy chain (Myh)* transcripts as distributed among all 11 mouse skeletal muscle tissues is plotted as a bar graph. The y-axis describes the percent of reads aligning to any given *Myh* transcript relative to all Myh-aligning reads. (B) Absolute expression of *Myh* transcripts among all 11 mouse skeletal muscle tissues is represented as a heat map. Absolute levels were plotted as a heat map to illustrate the dynamic range of *Myh* expression in muscle. Tissues are clustered by overall *Myh* similarity (top dendrogram). Notably, clustering is similar to, but distinct from, clustering done on global transcriptional profiles (**Figure 1D**). (C) The expression of *Myostatin (Mstn),* a gene specifically expressed by Type IIb (fast) fibers (10), is plotted versus *Myh4* expression. Each dot represents one of 11 skeletal muscle tissues (R^2^ = 0.473). (D) R-squared values for correlations with *Myh4* expression are plotted as a bar graph for 16 genes known to be specific for Type IIb fibers. Median R^2^ = 0.132; only one gene has an R^2^ > 0.5. (E) The expression of *Calsequestrin 2 (Casq2),* a gene specifically expressed by Type I (slow) fibers (10), is plotted versus *Myh7* epression. Each dot represents one of 11 skeletal muscle tissues (R^2^ = 0.412). (F) R-squared values for correlations with *Myh7* expression are plotted as a bar graph for 19 genes known to be specific for Type I fibers. Median R^2^ = 0.321; 7 of 19 genes (37%) tested show essentially no correlation with *Myh4* expression.

To explore these observations further, we made use of a single-fiber microarray study that identified genes enriched in slow (Type I) versus fast twitch (Type IIB) skeletal muscle fibers (10). These legacy data reveal that *Myostatin* (*Mstn*) is almost exclusively expressed in fast twitch fibers. Therefore, if fiber type composition determines gene expression patterns, we would expect close correlation between *Mstn* and *Myh4.* In actuality, there is only moderate correlation (R^2^ = 0.47, **Figure 2C**). Expanding on this result, we find weak correlation between genes enriched in fast twitch fibers and *Myh4* (median R^2^ = 0.132, **Figure 2D**). The low correlation between fiber type composition and gene expression signatures is also seen in slow twitch fibers (**Figure 2E** and **F**). Taken together, we conclude that fiber type based on *Myh* expression contributes to tissue-specific gene expression but is insufficient to establish the diversity of transcriptional patterns we observe.

Alternatively, developmental history may play an essential role in defining gene expression patterns in adult skeletal muscle. *Hox* genes are a family of transcription factors that establish anterior/posterior patterning and skeletal muscle specification during development (29). We therefore examined the expression of *Hox* genes in muscle tissues (**Figure 3A**). Clustering of tissues based on *Hox* gene expression reveals that the head and neck muscles cluster more closely with cardiac tissues than other skeletal muscles. Similarly, the diaphragm is more similar to the aorta than limb skeletal muscle. This is in marked contrast to clustering by the entire transcriptome, where each skeletal muscle is more similar to other skeletal muscles than any cardiac or smooth tissue. *Hox* gene expression in muscle recapitulates the developmental history of these tissues with respect to anterior/posterior axis formation. But, as these clusters are significantly different from those seen when comparing whole transcriptome expression patterns (**Figure 1D**), we conclude that *Hox* genes are insufficient to explain the mRNA diversity among adult skeletal muscle tissues.

**Figure 3:**
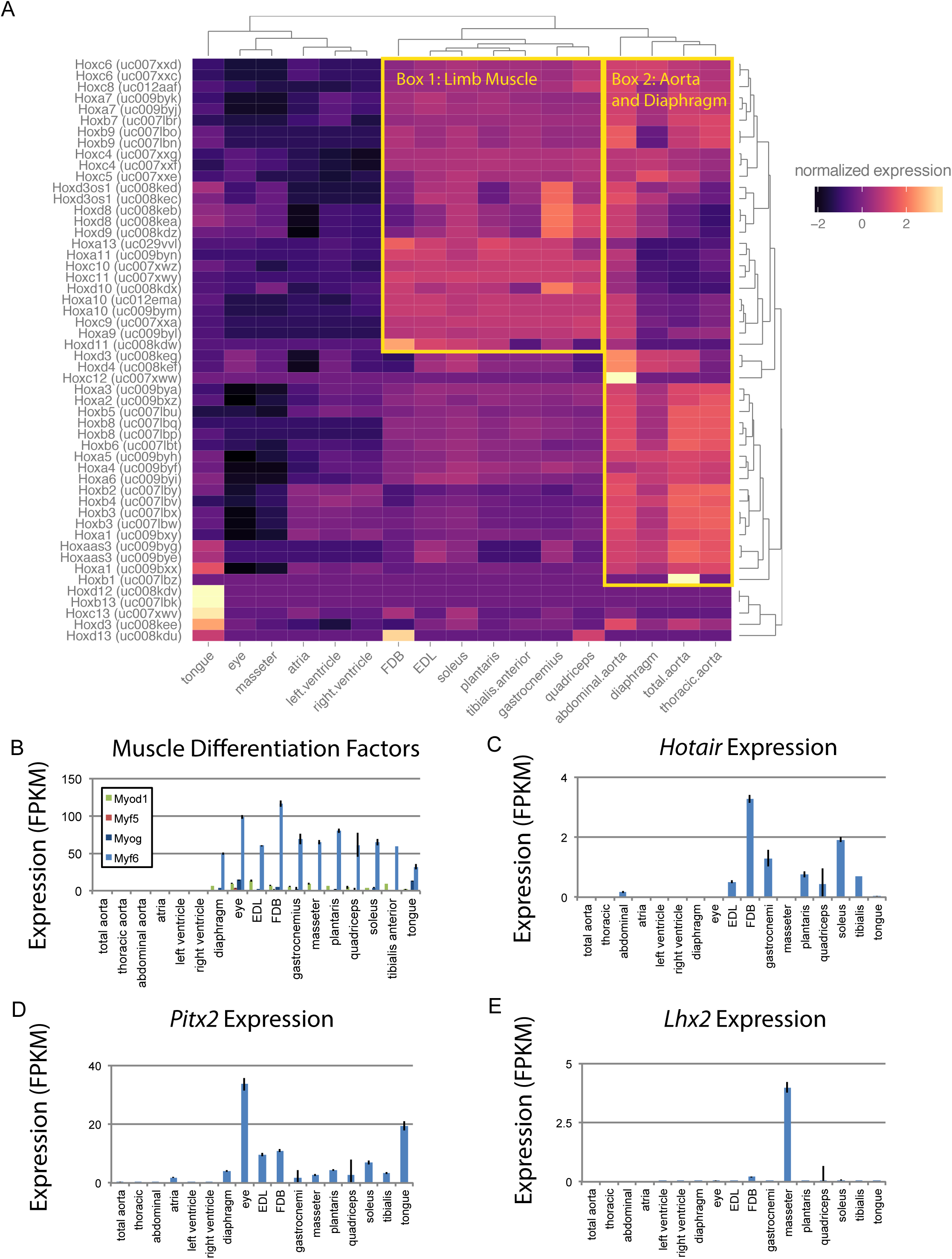
Developmental gene expression persists into adulthood in mouse skeletal muscle. (A) Normalized expression (Z-score by row) of all *Hox* gene transcripts is represented as a heat map. Row-normalization was chosen to display these data in a way that reveals the fine detail of all *Hox* genes, rather than those expressed at the highest levels. Overall similarity by *Hox* gene expression is represented as a dendrogram (top). One yellow box highlights a cluster of *Hox* genes expressed in limb skeletal muscle; a second yellow box highlights a cluster of *Hox* genes expressed in the aorta and diaphragm. (B) Expression of skeletal muscle differentiation factors *(Myod1, Myf5, Myog,* and *Myf6)* is shown as a bar graph. Skeletal muscle-specific expression of these genes persists into adulthood. (C) Expression of *Hotair,* a non-coding RNA involved in *Hox* gene regulation, is shown as a bar graph. *Hotair* expression is highly specific for a subset of skeletal muscle tissues involved in limb movement. (D) Expression of *Pitx2,* a gene involved in the development of head muscles, is shown as a bar graph. *Pitx2* expression is enriched in head and neck tissues such as the extraocular eye muscles and the tongue. (E) Expression of *Lhx2,* another gene involved in head and neck muscle development, is shown as a bar graph. *Lhx2* expression is highly specific for the *masseter*. Error bars are +/-S.E.M.

Similar to *Hox* family members, many developmentally significant genes maintain expression in adult muscle tissues. These include the myogenic regulatory factors *Myod1, Myf5, Myog,* and *Myf6,* which are expressed in, and exclusive to, skeletal muscle (**Figure 3B**). *Hotair* is a noncoding RNA that regulates *Hox* gene expression in development (30); as expected, it is expressed in all limb muscles (**Figure 3C**). In contrast, *Pitx2* is a transcription factor with a key role in driving extraocular muscle development (31), and *Lhx2* is a transcription factor important for *masseter* muscle development (32). The expression of both genes is consistent with regulatory activity that continues into adulthood and may contribute to maintaining proper mRNA expression patterns (**Figure 3D** and **E**). Notably, *Lhx2* is also involved in limb muscle development (33); it is of interest to determine how and why its expression is maintained in adult masseter but not adult limb tissues. We speculate that computational modeling of transcription factor networks may resolve the mechanisms underlying tissue-specific mRNA profiles, particularly since neither developmental history nor fiber type composition are sufficient to explain the diversity of the skeletal muscle transcriptome.

In fact, there are a considerable number of transcripts expressed in an entirely tissue-specific fashion in skeletal muscle. Plotting tissue specificity versus average expression reveals a bimodal distribution among all transcripts (**Figure 4A** and **B**). The majority of transcripts in skeletal muscle are expressed evenly across most skeletal muscle tissues, consistent with the notion that there is a minimal set of genes required to form sarcomeres and generate contractile force. Nevertheless, a smaller set of transcripts (~5%) show nearly exclusive expression in a single skeletal muscle tissue. Manual examination of these tissue-specific genes reveals that many are found in head and neck muscles that have the most divergent developmental history of tissues in this study. Nonetheless, we found examples of tissue-specific genes in the diaphragm (**Figure 4C**) and the *FDB* (**Figure 4D**). These data could therefore be used to generate transgenic mice with tissue-specific genetic manipulations, allowing follow-up studies to explore the mechanisms underlying muscle specialization.

**Figure 4:**
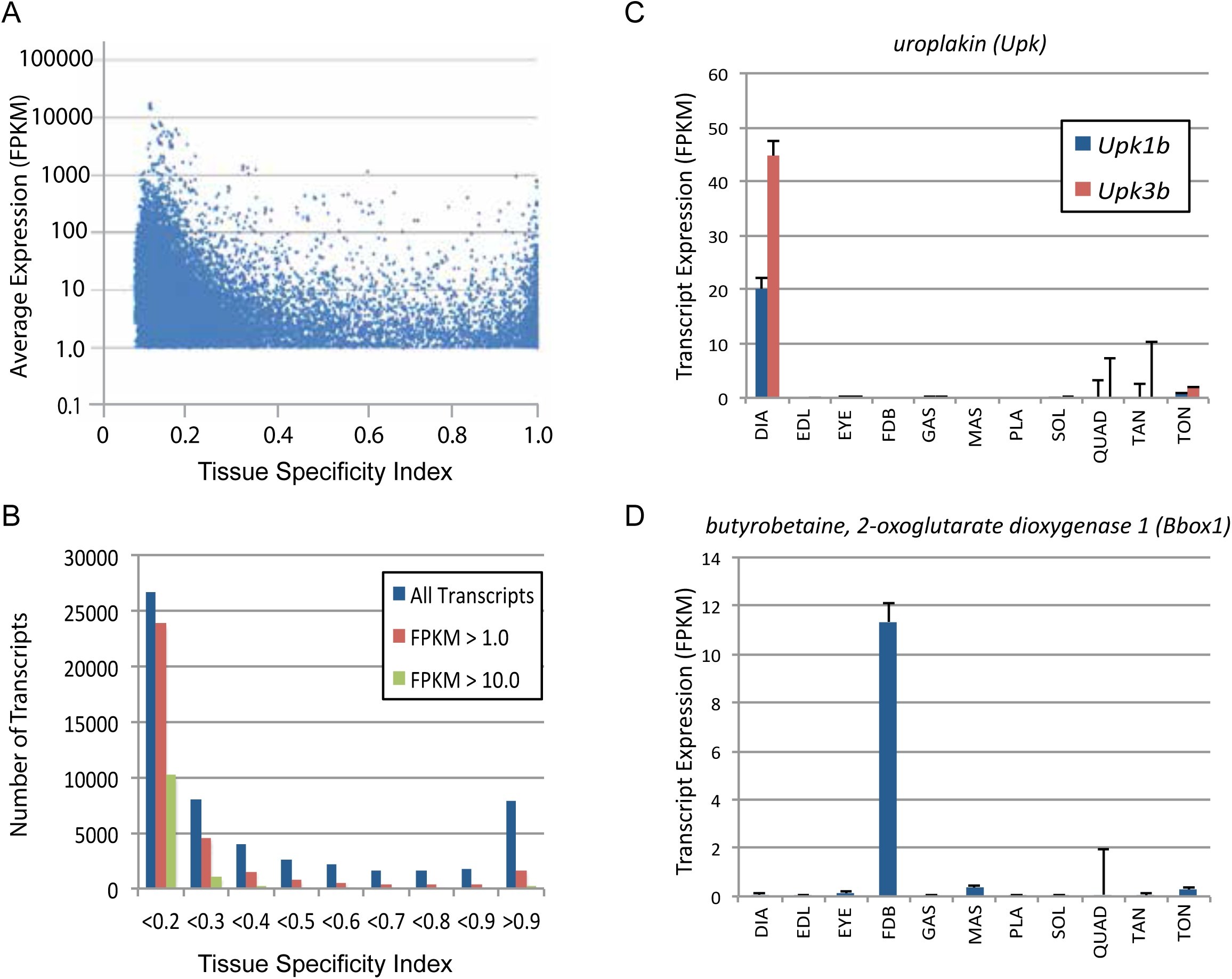
Subsets of genes are expressed specifically in different skeletal muscles. (A) Tissue specificity (maximum expression in *any* tissue divided by the sum total expression in *all* tissues) was calculated for all expressed transcripts in 11 different skeletal muscle tissues. The tissue specificity index ranges from 1.0 (completely specific for a single tissue) to 0.091 (evenly expressed among all 11 skeletal muscle tissues). Average expression is plotted versus tissue specificity for all transcripts expressed above an average FPKM of 1.0. The distribution of transcripts is quantified in (B), where the majority of expressed transcripts are expressed approximately equally across all 11 tissues (specificity index < 0.2), and a smaller group of transcripts are expressed with high specificity (specificity index > 0.9). Panel (C) shows two examples of genes *(Upk1b* and *Upk3b)* that are specifically expressed in the diaphragm. Panel (D) shows an example of a gene *(Bbox1)* expressed specifically in the *FDB* (Error bars are +/-S.E.M.).

These expression data reveal that the skeletal muscle transcriptome is considerably more complex than previously appreciated. Moreover, the unprecedented depth of sequencing coverage in muscle permits the discovery of heretofore-unannotated transcripts. Manual examination of “gapped reads” spanning two or more loci in the genome indicates that hundreds to thousands of splicing events occur in muscle that have not been previously described (**Figure S7A**). The majority of these novel splicing events result in either inclusion of novel exons or the exclusion of exons previously thought to be constitutive (**Figure S7B**). To validate this observation, we designed oligonucleotide primers (**Figure S7C**) specific for two novel exons of *Myosin light chain kinase 4 (Mylk4),* a gene with high expression (>200 FPKM) in many skeletal muscle tissues. PCR amplification and molecular cloning confirmed that these putative exons are included in full-length *Mylk4* transcripts in two different skeletal muscle tissues, *EDL* and *soleus* (**Figure S7D**). As the canonical splicing event linking exons 2 and 3 is detected by RNA-seq in every muscle sample in this study, albeit at levels below the threshold for detection in RT-PCR, we conclude that these putative exons are in actuality part of the predominant species of *Mylk4* mRNA.

There are significant differences in fiber type composition between analogous muscle tissues of different mammals. For example, mouse *soleus* is a mixture of fast-and slow fibers (**Figure 2A**), while rat *soleus* is almost entirely slow twitch (**Figure 5A**). Because of this, we asked whether gene expression differences in mice are conserved across species. Expression of orthologous genes in mouse versus rat tissues has moderate to high levels of correlation (R^2^ > 0.6, **Figure 5B**), despite the difference in fiber type composition noted above. This underscores the observation that tissue identity, rather than fiber-type composition, drives transcriptome diversity in muscle. Moreover, the vast majority of genes differentially expressed between *EDL* and *soleus* in both mice and rats changed in the same direction (**Figure 5C**). Taking this observation one step further, the fold change of all genes differentially expressed in mice *EDL* compared to *soleus* (**Figure 5D**) is largely consistent with the fold change of same genes in rat *EDL* versus *soleus* (**Figure 5E**). Sex differences did not dramatically influence differential gene expression between *EDL* and *soleus* (**Figure S8**). Fewer than 3% of transcripts were differentially expressed between male and female rat *EDL* (2.7%) and male and female rat *soleus* (1.9%). Of these differentially expressed genes, most are up-regulated in males. This observation agrees with previous studies (34) and is consistent with the possibility that androgen response elements influence sex-specific gene expression differences in skeletal muscle. These results indicate that differentially expressed genes are largely conserved between mice and rats and suggest that these data may predict gene expression in related species.

**Figure 5:**
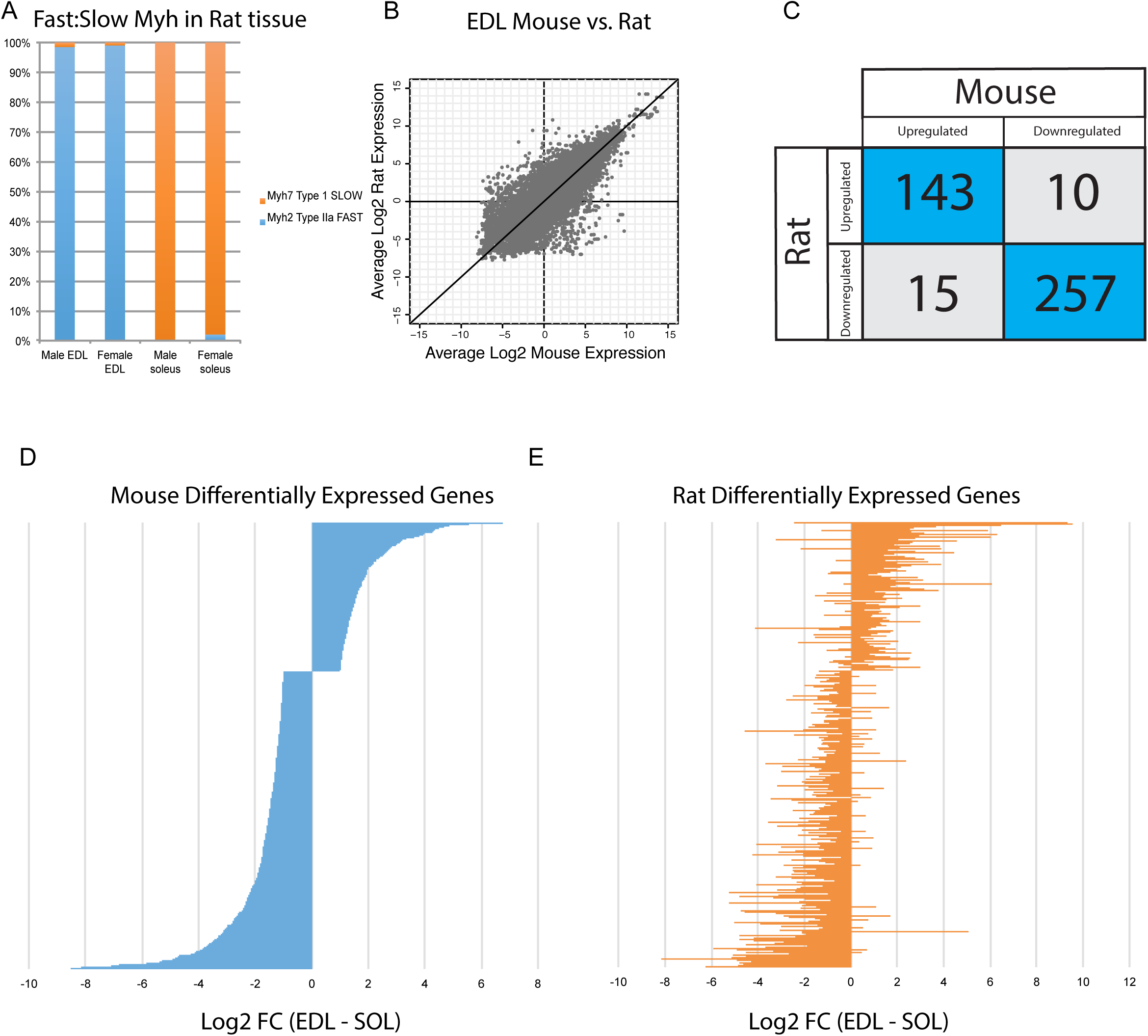
Differential gene expression is conserved between mice and rats. (A) The relative abundance of fast:slow *Myosin heavy chain (Myh)* transcripts in rat male and female *EDL* and *soleus* is shown as a bar graph. As expected, the fiber type composition of rat muscle is more homogeneous than mouse (compare with **Figure 2B**) (B) Scatter plot showing the overall similarity between mouse and rat transcriptomes in *EDL* (R^2^ = 0.637). Each dot represents a single orthologous gene shared between mice and rats. Correlation between the transcriptomes of mouse and rat *soleus* is essentially the same as for *EDL* described above (R^2^ = 0.662). (C) Of the 450 genes differentially expressed between *EDL* and *soleus* in both mice and rats, the majority (94%) were differentially expressed in the same direction (i.e., up-regulated in both mice and rats or down-regulated in both mice and rats). (D) The fold change for all rank-ordered differentially expressed genes in mice between EDL and soleus are plotted as a bar graph (N = 691, q < 0.05, fold change > 2). (E) The fold change *(EDL/soleus)* for the rat orthologues of the genes in (D) are plotted as a bar graph. The order of genes in (D) and (E) is identical. The majority of genes in rat (> 90%) show differential expression in the same direction as seen in mice (i.e., up-regulated in both mice and rats or down-regulated in both mice and rats).

Taken as a whole, there is considerable variance in gene expression profiles among skeletal muscle tissues, in stark disagreement with the assumptions of previous gene expression atlases. In the remainder of this paper, we will explore several vignettes illustrating the utility of these data as resource for generating testable hypotheses.

Based on the likely conservation of gene expression patterns in humans, we speculate that genes associated with disease and those encoding drug targets will be of particular importance to follow-up studies. Of the roughly 23,000 genes encoded by the mouse genome, over 50% are differentially expressed among mouse skeletal muscle tissues. Of these, 3,370 differentially expressed genes have human orthologs associated with disease, and 556 of those encode molecular targets of drugs on the market today (**Figure 6A**). These genes may contribute to the molecular mechanisms underlying differential disease susceptibility and pharmaceutical sensitivity in skeletal muscle tissues. As a resource for investigators, we provide a list of all differentially expressed genes specifically involved in skeletal muscle disorders (**Table S2**).

**Figure 6:**
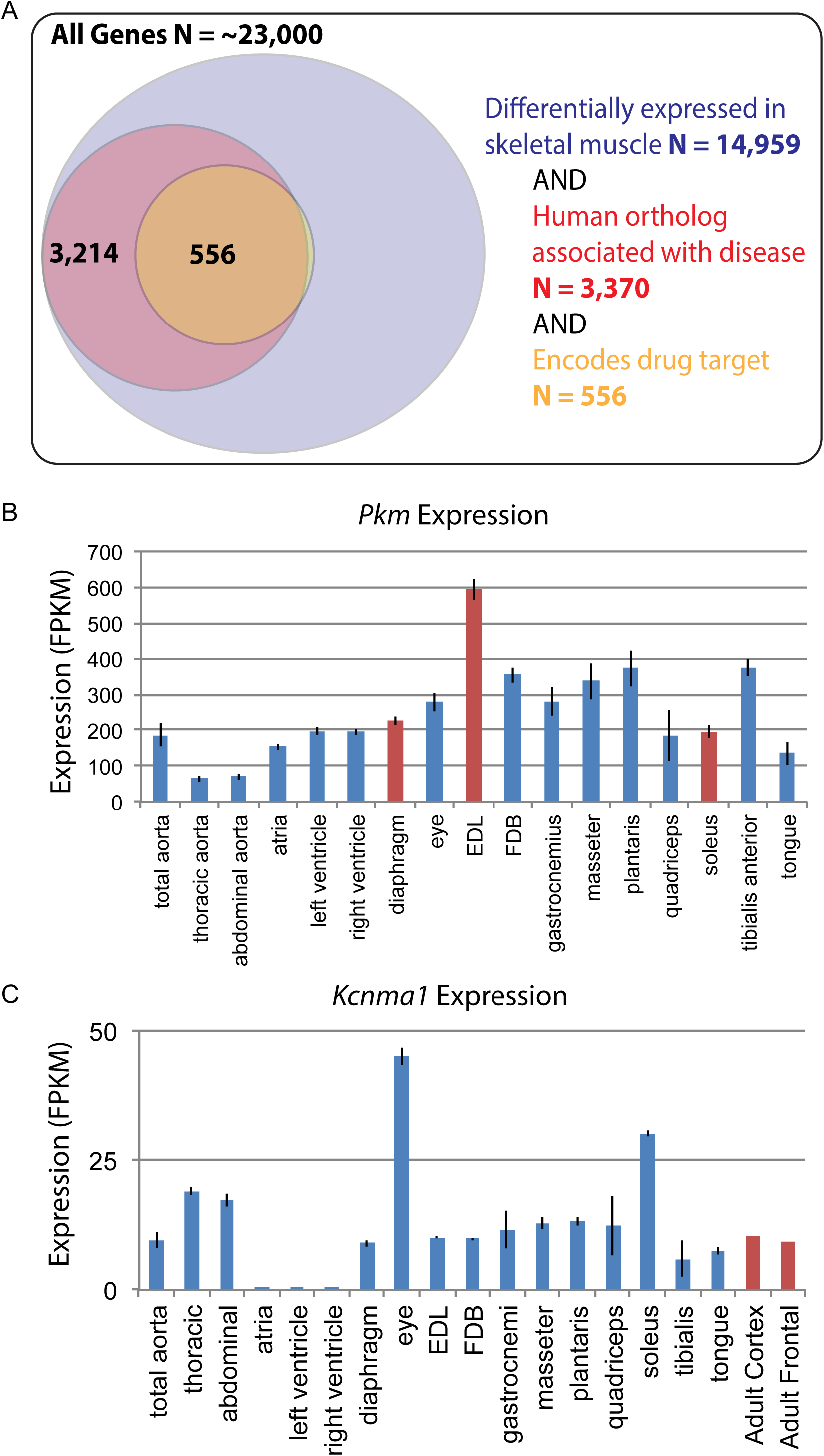
Differential gene expression in skeletal muscle may influence pathology and pharmacology. (A) Of roughly 23,000 genes in mice, 14,959 (~65%) are differentially expressed among skeletal muscle tissues (blue circle). Of these, 3,370 (~15% of all genes) have orthologs known to influence human disease (red circle). Of these, 556 (> 2% of all genes) encode the target of a marketed drug (orange circle). A small number of genes (N=13) encode a drug target and are differentially expressed in skeletal muscle, but are not annotated as being associated with human disease, which explains why a sliver of the Venn diagram does not entirely overlap between red and orange circles. (B) The expression of *Pkm,* a key gene involved in the pathogenesis of myotonic dystrophy, is shown as a bar graph. Data points highlighted in red *(EDL, soleus,* and diaphragm) represent tissues that have been empirically tested for sensitivity to degeneration in a mouse model of myotonic dystrophy (36). (C) Expression of *Kcnma1* is shown as a bar graph. *Kcnma1* encodes a potassium channel that is the molecular target of Chlorzoxazone, a drug prescribed as a muscle relaxant. Data points highlighted in red represent FPKM normalized gene expression in the adult cortex and adult frontal lobe as measured by the ENCODE consortium (38). Error bars are +/-S.E.M., when available.

To give one example of differential disease susceptibility, the aberrant expression of an embryonic isoform of *pyruvate kinase* (*Pkm*) is involved in the mechanism of myotonic dystrophy (35). Our data show that isoforms of *Pkm* are up-regulated by several standard deviations in *EDL* compared to all other muscle groups (**Figure 6B**). Myotonic dystrophy disrupts normal splicing and pathologically elevates *Pkm,* which in turn disrupts normal metabolism, decreasing oxygen consumption and increasing glucose consumption. Based on these observations, elevated expression of *Pkm* in adult tissues is hypothesized to be a critical step in the pathology of myotonic dystrophy (35). Since *Pkm* is expressed dramatically higher in *EDL* compared to all other muscle types (**Figure 6B**), we predict that *EDL* would be more sensitive to degeneration than other muscles. As myotonic dystrophy most dramatically affects certain subsets of muscle tissues, these observations suggest testable hypotheses regarding the underlying mechanism of disease susceptibility. We note with great interest that in mouse models, *EDL* is considerably more susceptible to muscle weakness than either diaphragm or *soleus* (36).

In addition to disease susceptibility, these data may help explain differential drug sensitivity in muscle. To give one example, the drug Chlorzoxazone (brand name: Lorzone) is used to treat muscle spasms. It is thought to act on the central nervous system by regulating a potassium channel encoded by the gene *Kcnma1* (37). Although *Kcnma1* is expressed at high levels in the central nervous system (38), it is also found at comparable levels in most skeletal muscle tissues (**Figure 6C**). The two exceptions are extraocular eye muscles and the *soleus* that have greater than three-fold higher *Kcnma1* expression than other muscle tissues. The differential abundance of Chlorozoxazone’s target could impact either the efficacy of this drug or the severity of its side effects in different muscles. The expression in skeletal muscle of mRNAs encoding Chlorzoxazone’s protein target calls into question the assumption in the literature that this drug acts exclusively through the CNS. Moreover, given that some commonly prescribed drugs, such as glucocorticoids, cause muscle wasting in specific subsets of muscle tissues (39), we speculate that this data set will be a valuable resource for exploring the mechanisms underlying differential drug sensitivity among skeletal muscles.

Skeletal muscles are endocrine tissues, expressing numerous secreted cues that influence the physiology and metabolism throughout the body. Termed “myokines”, these signaling molecules have important impacts on human health and disease (40). The total number of myokines is currently unknown. Our data contribute to this field by comprehensively defining candidate myokines for future study (**Figure S9** and **Table S2**). Dozens of genes encoding secreted proteins are differentially expressed among skeletal muscles at relatively high levels (N = 42 unique genes, FPKM > 10). One notable example is *Vegfa* (q-value ~ 10^-30^), a gene involved in angiogenesis, cardiac disease, cancer progression, and many other normal and pathological processes (41). Average expression of its predominant isoform is in the top 98^th^ percentile of all transcripts expressed by skeletal muscle. Moreover, the serum concentration of VEGF-A in normal healthy adults is considerably greater than serum levels of IL6, a canonical myokine (40). These data suggest that *Vegfa* expression in skeletal muscle has endocrine as well as paracrine functions. Moreover, maximal expression of *Vegfa* in the diaphragm is nearly 10-fold greater than its minimal expression in the FDB, indicating that there are muscle-specific mechanisms for regulating *Vegfa* levels in particular and myokine levels in general.

Regenerative medicine has made considerable progress in generating skeletal muscle from stem cells (42). Nevertheless, engineered tissues have important deficiencies in generating sufficient force and forming appropriate neuromuscular synapses (43). Given the extensive diversity observed among skeletal muscle tissues, we speculate that the inability to form proper synaptic connections in engineered tissue may be due to the expression of inappropriate or incomplete transcriptional programs. In other words, differentiating stem cells into generic skeletal muscle may not recapitulate all the cues necessary for muscle-specific synapse formation. Therefore, we examined the differential expression of genes implicated in synapse assembly (**Figure S10**). Dozens of transcripts involved in synapse formation are expressed at relatively high levels (FPKM > 10), suggesting that they are bonafide skeletal muscle mRNAs, rather than contamination from nearby neurons. One illustrative example, *Fbxo45*, is an E3 ligase involved in synapse formation. Mouse knock-outs of *Fbxo45* have disrupted neuromuscular junction (NMJ) formation in the diaphragm, resulting in early lethality (44). Moreover, the C. elegans ortholog, *FSN-1,* also disrupts NMJ formation, with some synapses being over-developed while others are under-developed (45). We speculate that this protein and its orthologs may play a conserved role in tissue-specific NMJ assembly.

## Discussion

General textbook discussions of skeletal muscle typically focus on developmental patterning, neuromuscular synapse physiology, or the biophysics of contractile properties (46). This reflects the generally held belief that adult skeletal muscle is interesting only as a mechanical output of the nervous system. Functional genomics studies have acted on this assumption to the extent that every gene expression atlas generated to date has selected at most one skeletal muscle as representative of the entire family of tissues. As such, the null hypothesis of this study was that gene expression profiles would be largely similar among skeletal muscle tissues.

However, as more than 50% of transcripts are differentially expressed among skeletal muscles (**Figure 1C**), and 13% of transcripts are differentially expressed between any two skeletal muscle tissues on average (**Figure 1E**), the data are entirely inconsistent with the null hypothesis. These results indicate that there is no such thing as a representative skeletal muscle tissue. Instead, skeletal muscle should be viewed as a family of related tissues with a common contractile function, but widely divergent physiology, metabolism, morphology, and developmental history. Based on these antecedents, it should come as no surprise that the transcriptional programs maintaining skeletal muscle specialization in adults are highly divergent as well.

This study is the first systematic examination of transcriptome diversity in skeletal muscle. At greater than 200 million aligned short nucleotide reads per tissue and six biological replicates apiece, this data set is unprecedented in its accuracy and reproducibility (see **Figures 3, S2, S4,** and **S6**). Moreover, the depth of sequencing allows the detection of previously unannotated transcripts that may play a role in muscle physiology (see **Figures S3** and **S7**).

Besides establishing that skeletal muscles have considerable differences in their transcriptomes, we believe the key significance of this paper will be as a resource for future studies. Therefore, we have made our analyzed data freely available (http://muscledb.org), and all raw data may be downloaded from NCBI’s GEO. This resource will allow investigators to perform analyses beyond the scope of this paper, including pairwise Gene Ontology analyses of differentially regulated genes and pathways. Likewise, these data may be mined to generate muscle-specific *Cre-*recombinase mouse strains for genetically manipulating specific muscle groups. Most importantly, these data will provide the foundation for computational modeling of transcription factor networks, a method we believe will uncover the genetic mechanisms that establish and maintain muscle specialization.

We acknowledge that many processes besides steady-state mRNA levels regulate protein expression. Nevertheless, regulation of mRNA expression is unquestionably of biological importance in muscle cells, and transcriptional profiling predicts the majority of protein expression levels even in highly dynamic settings (47). Moreover, the larger dynamic range of mRNA measurements permits a more comprehensive description of expression profiles than would be possible with existing proteomic technology. We look forward to proteomic studies making use of these mRNA data, especially the identification of novel spliceforms, to generate improved catalogs of protein expression in skeletal muscle.

We further acknowledge that the samples collected and analyzed in this study are bulk tissues rather than single-fiber preparations, and that contaminating tissues such as vasculature and immune cells may influence some gene expression measurements. However, we see agreement between our data and the few single-fiber profiling papers in the literature (10), indicating that this potential bias is unlikely to confound the major observations herein. Furthermore, we emphasize that the majority of studies in the field use bulk tissues rather than single cell preparations. As such, our experimental design yields the greatest possible consistency with previous and future studies. In short, we believe whole tissue expression profiling provides the rationale and a critical reference point for follow-up work examining single-fiber gene expression.

Acquired and genetic diseases show remarkable selectivity in which muscles they affect and which muscles they spare. For example, Duchenne muscular dystrophy severely affects the diaphragm and proximal limb extensors, while oculopharyngeal dystrophy causes weakness in the neck, facial, and extraocular muscles (17). At present, there is no satisfying explanation for how this occurs. A reasonable hypothesis is that intrinsic properties of muscle cells, such as gene expression, determine their sensitivity to different pathological mechanisms. These data are a starting point for future studies on how specialized transcriptional programs in muscles are maintained and how they ultimately influence disease. As an illustrative example, we note that the naturally elevated expression of *Pkm* in *EDL* may in part explain how this muscle is most dramatically affected in mouse models of myotonic dystrophy (**Figure 6**). Related to this, the use of engineered skeletal muscles in regenerative medicine has been limited by poor contractile properties and inadequate neuromuscular synapse formation (43). We speculate that this deficiency may be corrected if engineered tissues are programed to express appropriate synaptic assembly proteins (**Figure S10**).

Finally, skeletal muscle is an endocrine organ that regulates many normal and pathological processes, including sleep, bone health, diabetes, cancer, and cardiovascular disease (48–50). At roughly 40% of an adult human’s body weight, skeletal muscle has an enormous capacity to influence other tissues through the expression of local or systemic signaling molecules. This study reveals extensive differential expression of putative myokines with largely unexplored functional significance (**Figure S9**). As such, we predict these data will be instrumental in future studies of the endocrine mechanisms through which skeletal muscle regulates health and disease.

## Methods

### Animal care and tissue collection

Adult male *C57Bl6J* mice were acquired from Jackson Laboratories at ten weeks of age. They were housed in light tight cages with a 12L:12D light schedule for four weeks with water and normal chow *ad libitum.* At 14 weeks of age, mice were sacrificed at between two and five hours after lights-on (i.e. ZT 2-5), and muscle tissues were rapidly dissected and flash frozen in liquid nitrogen for subsequent purification of total RNA. Adult male and female *Sprague Dawley* rats were obtained from Charles River (Wilmington, MA) at 12 weeks of age. Rats were housed in pairs with a 12-h:12-h light/dark cycle, and standard rat chow and water were provided ad libitum. At 14 weeks of age, rats were sacrificed at between two and five hours after lights-on (i.e. ZT 2-5), and muscle tissues were rapidly dissected and flash frozen in liquid nitrogen for subsequent purification of total RNA. Three animals were sacrificed per biological replicate. All animal procedures were conducted in compliance with the guidelines of the Association for Assessment and Accreditation of Laboratory Animal Care (AAALAC) and were approved by the Institutional Animal Care and Use Committee at University of Kentucky.

### RNA purification and library preparation

Between 5-20 mg of frozen tissue were manually homogenized in Trizol reagent (Invitrogen), and total RNA was purified using a standard chloroform extraction. RNA samples for gene expression analysis were mixed with an equal volume of 70% ethanol and further purified with RNEasy columns (Qiagen) using the manufacturer’s protocol. RNA was purified from tissue collected from individual mice; samples from three individual mice were then pooled together in equimolar amounts for further analysis.

To assess RNA integrity, aliquots of each sample were denatured for 2 minutes at 70°C and analyzed on the Agilent 2100 Bioanalyzer using Eukaryote Total RNA Nano chips according to manufacturer’s protocol. RNA integrity numbers (RINs) for all samples were above 8.0 with a median RIN of 9.2. Libraries were prepared using the Illumina Truseq Stranded mRNA LT kit using single end indexes according to manufacturer’s protocol. Approximately 500 ng of total RNA was used as starting material and amplified with 13 cycles of PCR. Libraries were validated for size and purity on the Agilent 2100 Bioanalyzer using DNA 1000 chips according to manufacturer’s protocol.

### RNA-sequencing and analysis

Pilot runs to verify library integrity were sequenced on an Illumina MiSeq (University of Missouri—St. Louis), and subsequent sequencing was performed on an Illumina HiSeq 2500 (University of Michigan) or HiSeq 3000 (Washington University in St. Louis). All sequenced reads were 50bp, single-end. Raw reads were aligned to the genome and transcriptome of *Mus musculus* (build mm10) or *Rattus norvegius* (build rn5) using RNA-seq Unified Mapper (RUM) (51) with the following parameters: *“--strand-specific --variable-length-reads --bowtie-nu-limit 10 --nu-limit 10”*. Mouse and rat gene models for RUM alignments were based on UCSC gene models updated as of December 2014. 65%-92% of reads uniquely mapped to the genome/transcriptome, and the total number of aligned reads (including unique and non-uniquely aligned reads) was at least 94.7% for every replicate sample (**Table S1**). FPKM values for each transcript/exon/intron were calculated by RUM using strand-specific unique reads normalized to the total number of reads uniquely aligned to the nuclear genome. To account for variability in the mitochondrial content of different muscles, uniquely aligned mitochondrial reads were excluded from the denominator. R-squared values of log-transformed FPKMs between biological replicates of the same tissue were generally ~;0.94 (**Figure S2**), and internal controls (e.g. **Figure S4**) were used to verify the biological validity of these measurements.

Differential expression between tissues was determined by one-way ANOVA of log-transformed FPKM values and adjusted for multiple testing using a Benjamini-Hochberg q-value (52). Unless otherwise noted, a q-value less than 0.01 and fold-change greater than 2.0 among transcripts expressed with an FPKM > 1.0 was deemed statistically significant. Transcripts were deemed to be expressed at FPKM > 1.0, unless otherwise noted. Disease and drug target associations were identified using public data sets (DrugBank v5.0.9 and DisGeNET or Ingenuity Pathway Analysis v01-12). Mouse genes were mapped to human orthologs using the BiomaRt package for R (53). Dendrograms were produced in R. A distance matrix was calculated for all (**Figure 1D**) or subsets (**Figure 2B, 3A**) of transcripts using the `<u>dist</u>` function using the standard options (Euclidean distance; **Figure S5**). Hierarchical clustering was performed using the `<u>hclust</u>` function using the complete agglomeration method. The code used to produce the heatmaps and dendrograms is freely available on <u>GitHub</u>.

### Validation and novel splicing events

Independent biological replicates of mouse *EDL* and *soleus* tissues were collected as described above. Total RNA was purified and integrity was verified as described above. Reverse transcription reactions were performed with 500ng starting material using the manufacturer’s protocol (TaqMan Fast Universal PCR Master Mix, Applied Biosystems). qPCR was performed with ABI Taqman probes on a Stratagene MX3005 instrument (Pcp4l1:mm01295270_m1, Fam129a:mm00452065_m1, Fhl2, mm00515781_m1, Tsga10:mm01228282_m1, Stau2:mm00491782_m1, Prkag3:mm00463997_m1, Mstn:mm01254559_m1, Plcd4:mm00455768_m1, Myl1:mm00659043_m1, Igfbp5:mm00516037_m1, Ipo8:mm01255158_m1) using the manufacturer’s recommendations.

To identify novel splicing events, previously unknown introns were identified from the junctions_all.rum file in RUM’s output. These results were then sorted by the number of reads aligning to that splice junction and manually curated for follow-up studies. To validate novel splicing events, PCR reactions were performed using 1ul of the reverse transcription reaction using the manufacturer’s protocol (Clontech Takara PCR kit). PCR products were visualized on a 1% agarose gel using conventional methods. The primary PCR products for *EDL* and *soleus* (**Figure S7D**) from primers Exon 2-> Exon 2.1 and Exon2 -> Exon3 were excised, purified, and TOPO cloned using the manufacturer’s protocol (TOPO TA, Thermo Fisher). Cloned fragments were sequenced using conventional Sanger methods. All PCR primers were ordered from IDT; primer locations are described in **Figure S7C**. Primer sequences as follows: Exon 2 – AGGATCTCAGATTTGCTCACG, Exon 2.1 – GGATCCACTTTCCAGAATGC, Exon 2.2 – CATCTTTGCACCTGCATTC, Exon 3 – TATGGTCCAACCGTGCACTA, CtrlF – AGTGTGGGCGTCATCACAT, CtrlR – GTGGAGCTTGTGGTCTGACA.

### Data Availability

All raw data are available on NCBI’s Gene Expression Omnibus (accession number: GSE100505), and transcript-level expression values can also be downloaded from MuscleDB (http://muscledb.org/), a web application built using the ExpressionDB platform (54). We note that all tracking software has been disabled for the duration of peer-review.

## Acknowledgements

We thank Jeanne Geskes, Robert Lyons and the University of Michigan DNA sequencing core facility for assistance with next-generation sequencing. We thank John Hogenesch (Cincinnati Children’s Hospital), Jeff Haspel (WashU Department of Medicine), Patty Parker (UMSL) and members of the Hughes and Esser laboratories for helpful discussion and technical support throughout this project. We thank Dr. Michael Chicoine (WashU Department of Neurosurgery) for inestimable contributions without which this paper would never have been written. Work in the Gong lab is supported by NIH award NIH HL106843. Work in the Esser lab is supported by NIH award R01AR066082. Work in the Hughes Lab is supported by NIH award R21AR069266 and start-up funds from the Department of Medicine at Washington University in St. Louis. We thank the Genome Technology Access Center in the Department of Genetics at Washington University School of Medicine for help with genomic analysis. The Center is partially supported by NCI Cancer Center Support Grant #P30 CA91842 to the Siteman Cancer Center and by ICTS/CTSA Grant# UL1 TR000448 from the National Center for Research Resources (NCRR), a component of the National Institutes of Health (NIH), and NIH Roadmap for Medical Research. This publication is solely the responsibility of the authors and does not necessarily represent the official view of NCRR or NIH.

---

**Figure S1:**
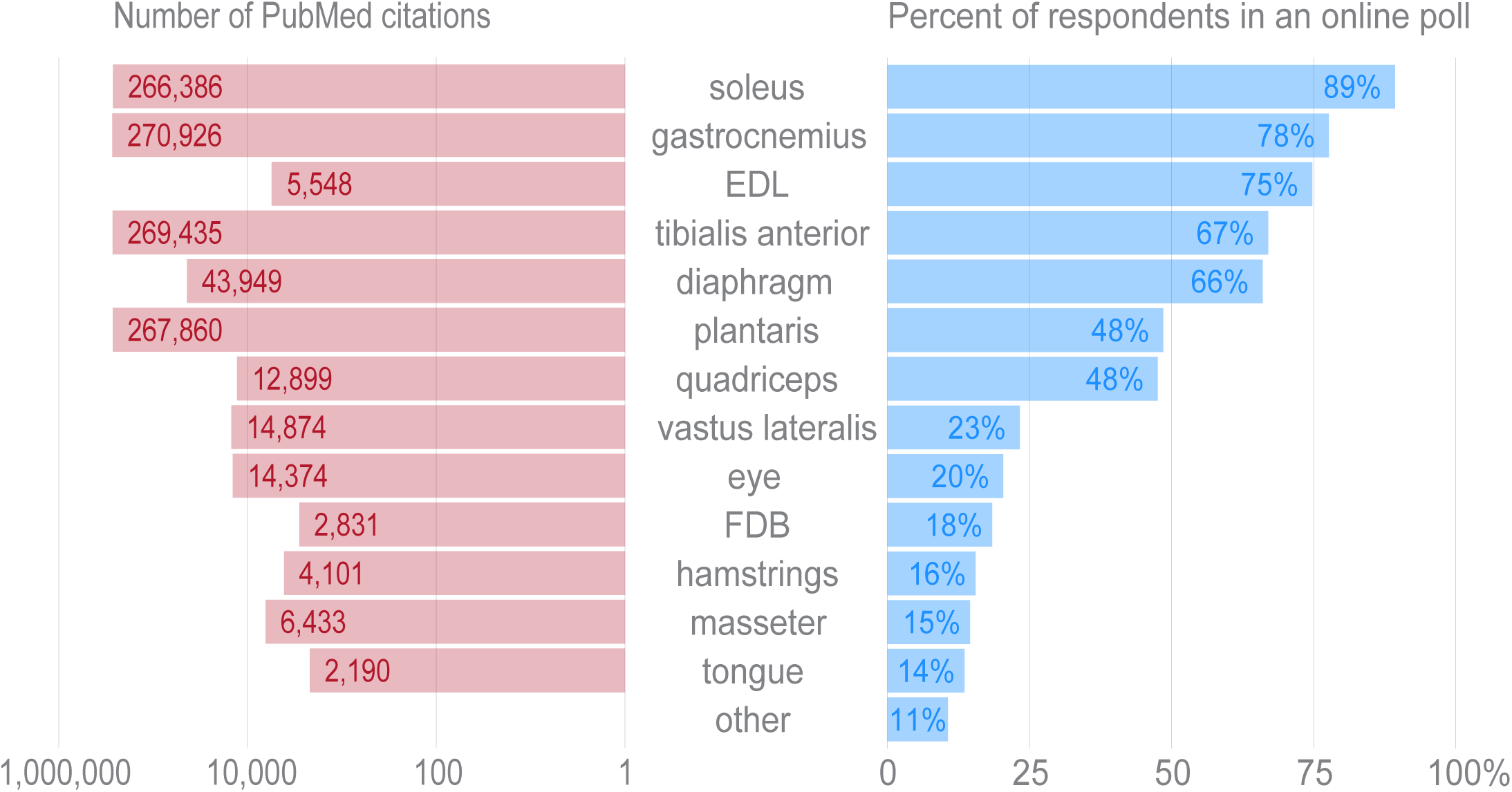
Tissues were selected for transcriptional profiling to maximize the utility of these data to the skeletal muscle field. In order to ascertain which tissues are of the greatest general interest to the skeletal muscle researchers, we distributed an online poll asking respondents to vote whether any given tissue was of interest to their laboratory (right panel). Over one hundred responses from principal investigators, graduate students, and postdocs in the field were recorded. The choice of “other” was ranked lower than any other tissue, and write-in responses showed no evidence of broad support for any tissues beyond those included here. *Quadriceps* were selected for profiling as they were favored in the poll over the more specific *vastus lateralis* by a ~2:1 ratio. *Masseter* and tongue were chosen for transcriptional profiling over hamstrings to increase the developmental and anatomical diversity of the tissues in this data set. In order to guard against selection bias for our online poll, we queried NCBI’s PubMed to identify the number of published papers on each muscle tissue (left panel). The overall distribution of PubMed citations mirrors the human poll.

**Figure S2:**
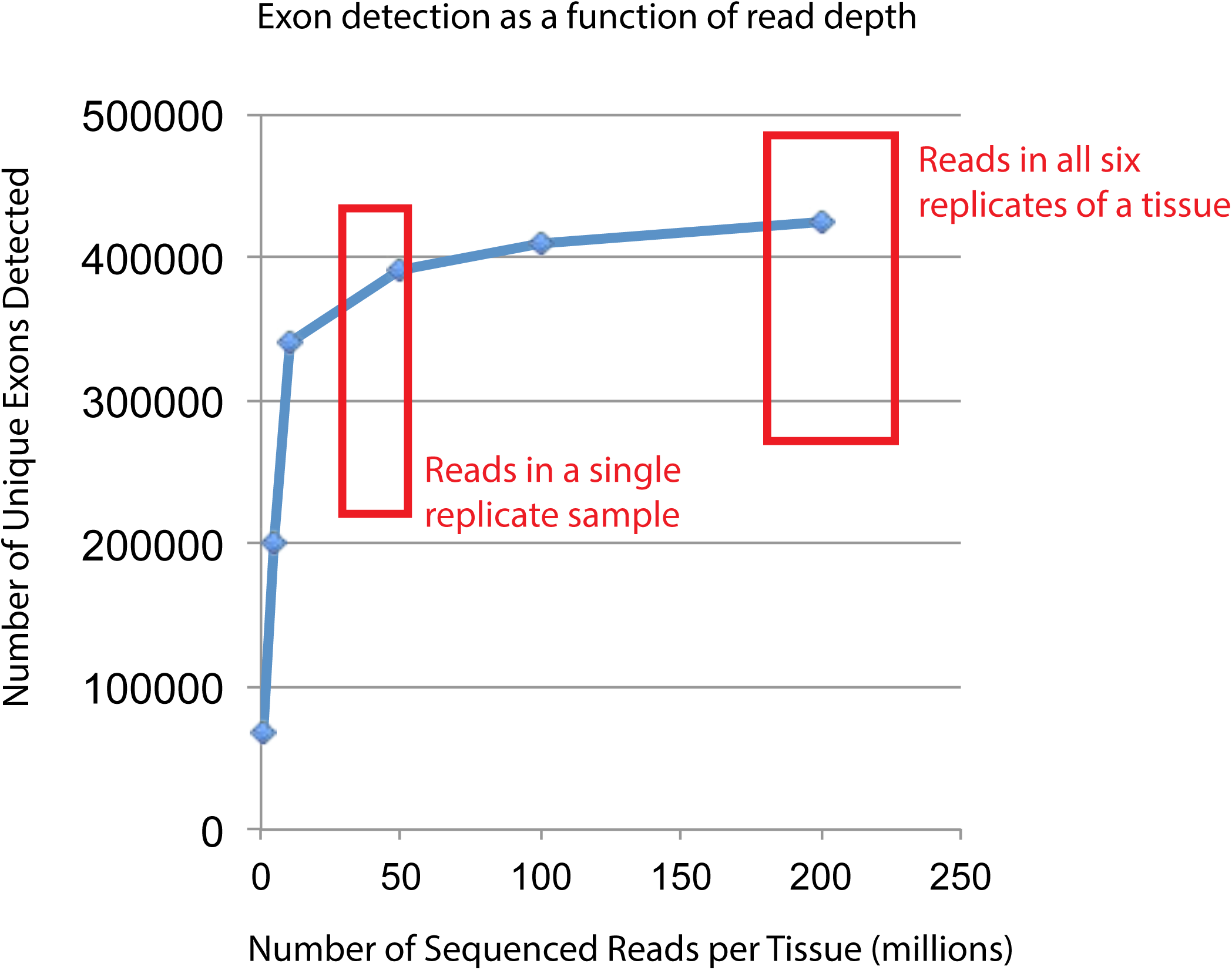
Empirical simulations show that the read depth of this study approaches saturation for detecting expressed exons. In order to verify that the RNA-seq read depth was sufficient to detect most expressed transcripts, one specific tissue (mouse *FDB)* was randomly down-sampled as previously described (55) to generate six different sequencing depths (200, 100, 50, 10, 5, and 1 million reads). These reads were aligned to the mouse genome and transcriptome using RUM (51), and uniquely aligned reads were used to identify putatively expressed exons. The number of unique exons detected as expressed (FPKM > 0) is plotted versus the total number of reads collected. The two red boxes indicate either the approximate number of reads for any given biological replicate or the approximate number of reads per tissue (summing all six replicates). Most single replicates have been sequenced to a depth that approaches diminishing returns for exon detection, while every tissue has been sequenced to a depth that guarantees that the vast majority of truly expressed exon are detected.

**Figure S3:**
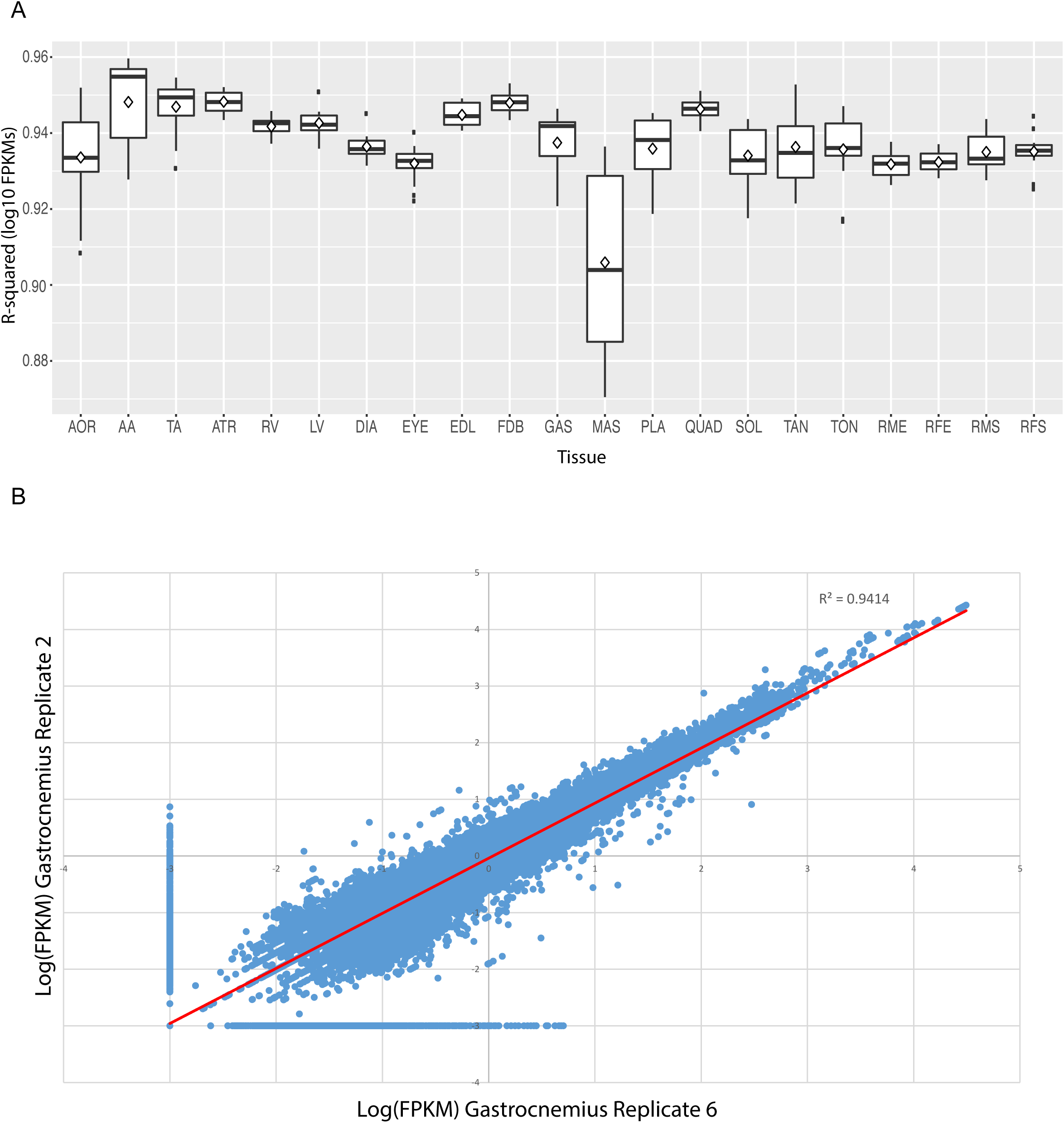
FPKM values show high reproducibility between replicate samples. (A) Box and whisker plots of R-squared values of pairwise comparisons between replicates of each tissue. Every median R-squared is greater than 0.90. Excluding *masseter,* every median R-squared is greater than 0.93. AOR = total aorta, AA = abdominal aorta, TA = thoracic aorta, ATR = atria, RV = right ventricle, LV = left ventricle, DIA = diaphragm, EYE = extraocular eye muscle, EDL = *extensor digitorum longus,* FDB = *flexor digitorum brevis,* GAS = *gastrocnemius,* MAS = *masseter,* PLA = *plantaris,* QUAD = *quadriceps,* SOL = *soleus,* TAN = *tibialis anterior,* TON = tongue, RME = male rat *extensor digitorum longus,* RFE = female rat *extensor digitorum longus,* RMS = male rat *soleus,* RFS = female rat *soleus*. (B) Scatter plot of all transcript-level FPKM values for two representative replicates (mouse *gastrocnemius* replicates 2 and 6). FPKM values are log10 transformed; any FPKM = 0 was plotted as an arbitrarily low expression value (FPKM = 0.001) to avoid log-transforming zero.

**Figure S4:**
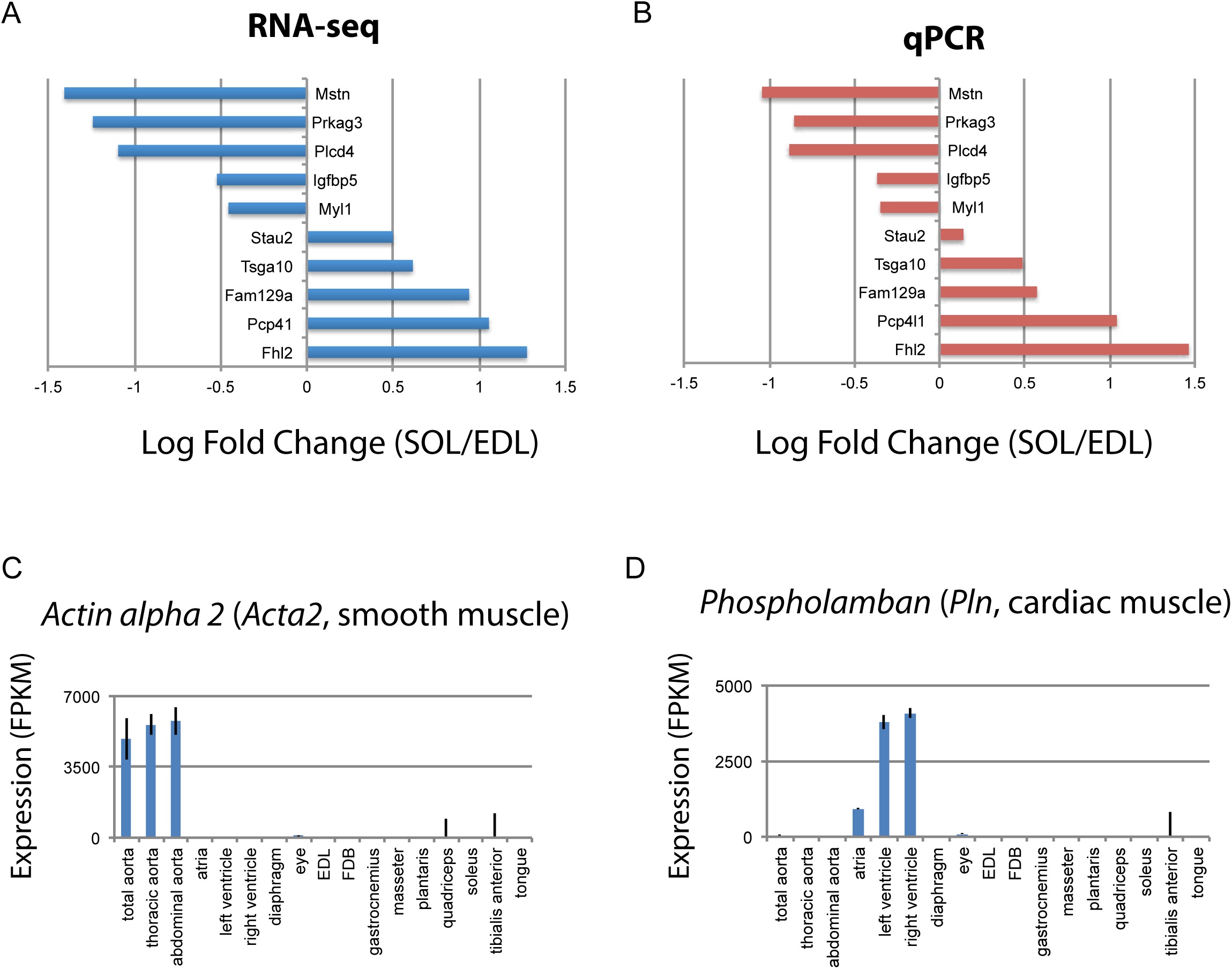
Internal controls demonstrate the reliability of differential expression analysis. 10 genes differentially expressed between mouse *EDL* and *soleus* were selected with log2 fold changes ranging from −1.5 to 1.5. Panel (A) shows the *EDL/soleus* log2 fold changes as a bar graph from RNA-seq data. Panel (B) shows the log2 fold changes for the same genes, normalized to the house-keeping gene *Importin 8*, measured by qPCR on biological samples *collected independently of those used for RNA-seq*. Internal controls based on known tissue-specific genes also corroborate the reliability of these data. (C) Expression of *Acta2,* a smooth muscle-specific gene, is plotted as a bar graph. (D) Expression of *Pln,* a cardiac muscle-specific gene, is plotted as a bar graph. Error bars are +/-S.E.M.

**Figure S5:**
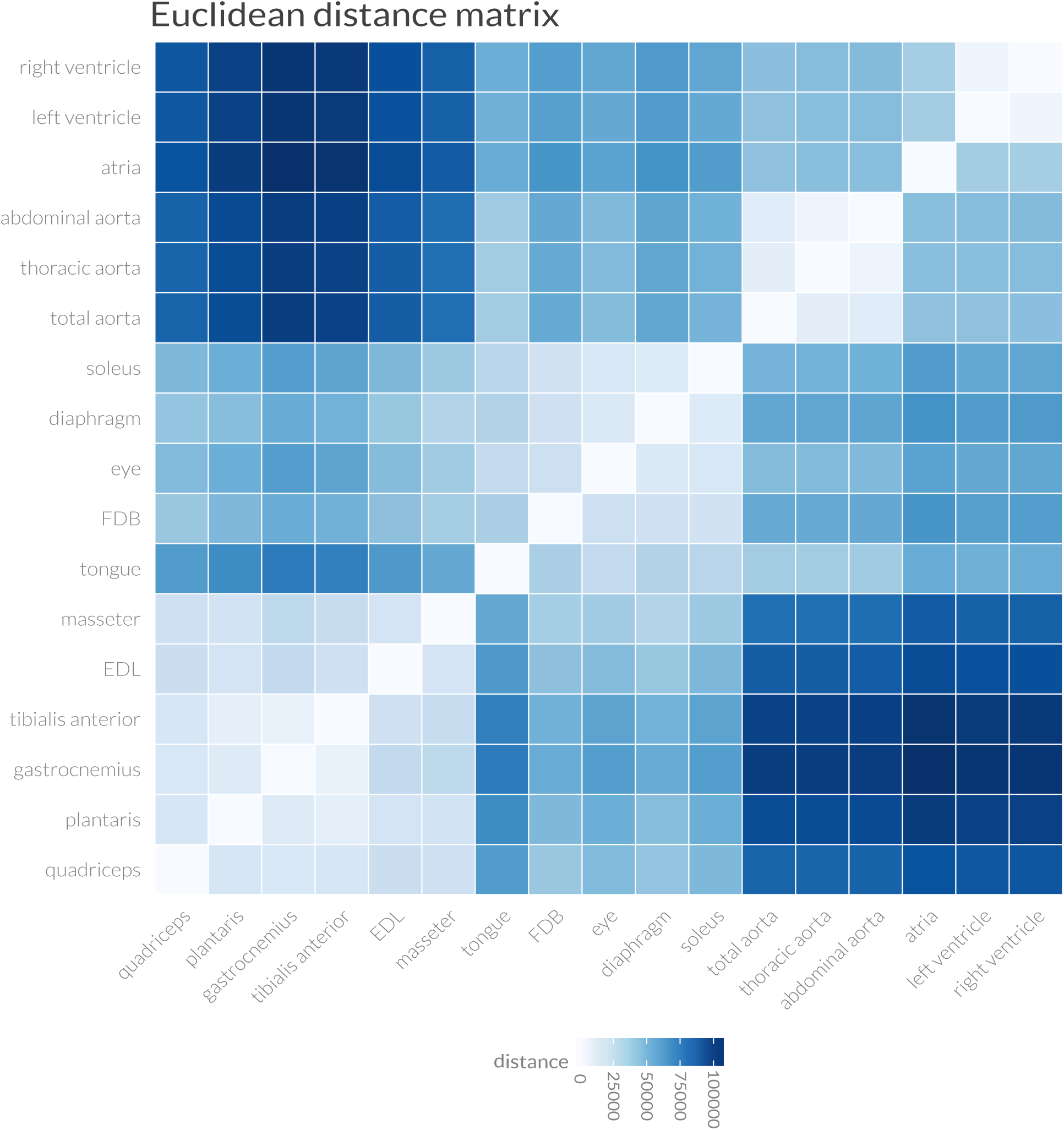
Euclidean distance measurements support the clustering of muscle transcriptomes into four distinct groups. Pairwise Euclidean distances between the entire transcriptome of every mouse tissue are shown as a heat map. Overall similarity reveals four distinct groupings: (1) cardiac muscle, (2) smooth muscle / aorta, (3) limb (except soleus) and *masseter* skeletal muscles, and (4) other skeletal muscles, including soleus. These data were used to generate the dendrogram shown in **Figure 1D**.

**Figure S6:**
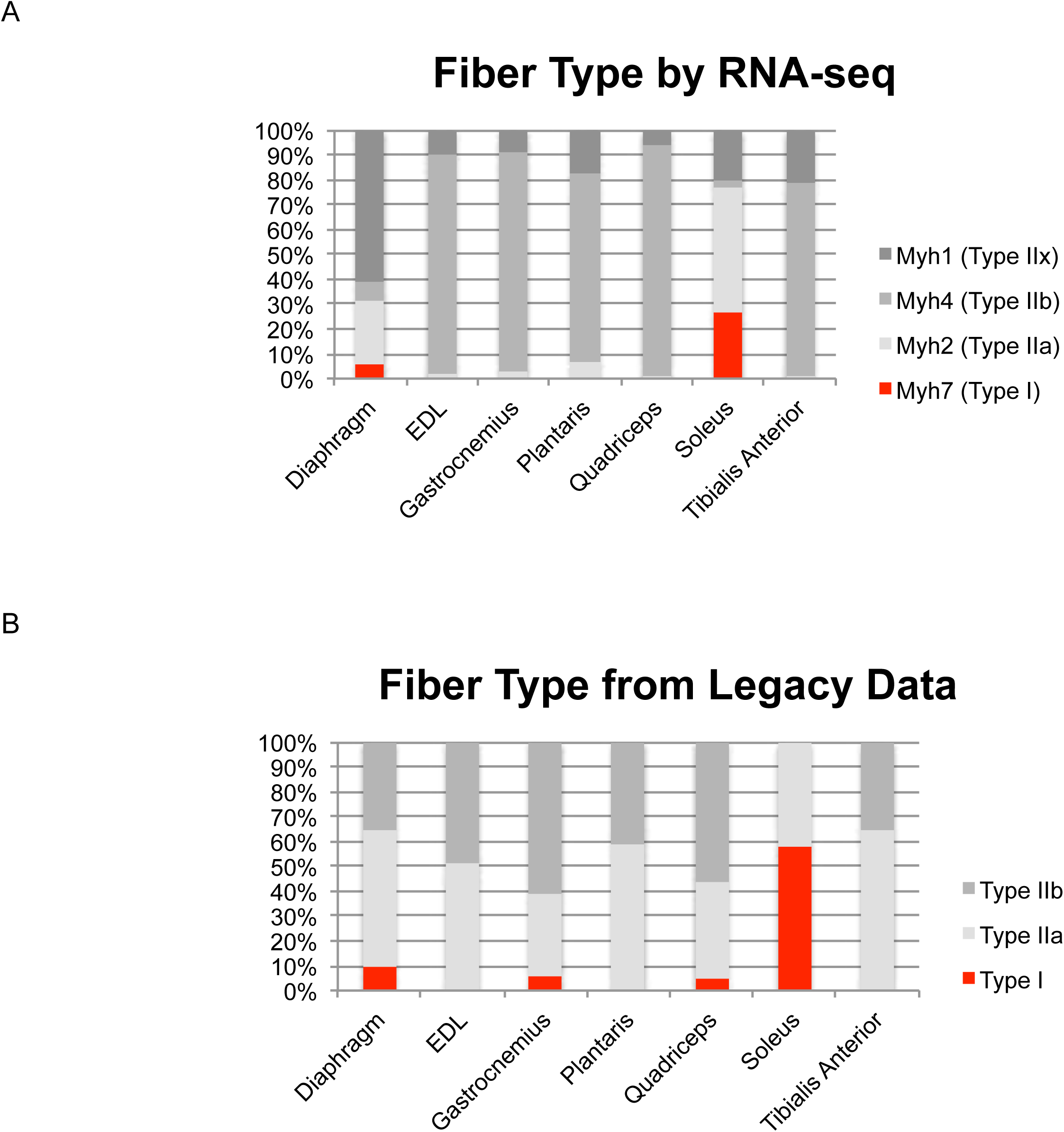
Identifying skeletal muscle fiber type by *Myh* expression agrees with legacy data. (A) Relative expression levels of the four principal *Myosin heavy chains (Myh1, 2, 4,* and 7) are shown as a bar graph in select mouse muscle tissues. Fast twitch myosins are shown in grey; slow twitch myosin is shown in red. (B) We identified legacy studies that characterized *Myh* expression in mouse diaphragm (56) as well as the *EDL, gastrocnemius, plantaris, quadriceps, soleus* and *tibialis anterior* (57); the relative compositions of fast/slow twitch fibers is represented as a bar graph. As above, fast twitch *myosins* are shown in grey; slow twitch myosins are shown in red. We note that the previous studies cited herein: (1) measured fast/slow composition in female as opposed to male mice, which is expected to increase the relative proportion of slow twitch fibers, (2) used histology, which is limited to analyzing cross-sectional area of serial sections, rather than the entire muscle body as in the present study, (3) used enzymatic methods to distinguish different fiber types, which limits the extent to which different fast twitch fibers can be resolved, notably neglecting Type IIx fibers. With these caveats in mind, to a first approximation, RNA-seq data of *Myh* expression agrees with legacy data of fiber type composition.

**Figure S7:**
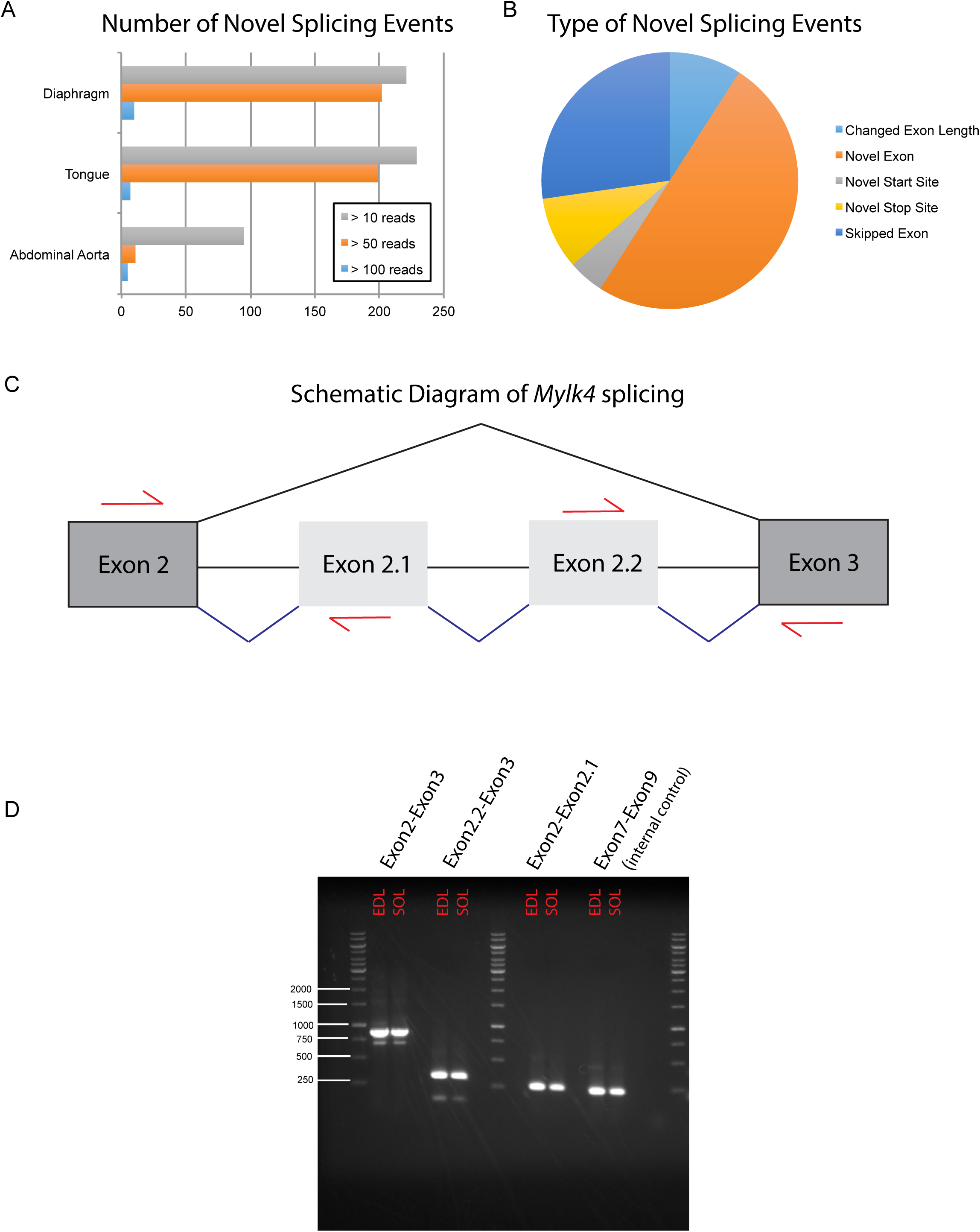
Deep sequencing the muscle transcriptome reveals numerous novel splicing events. In order to detect previously unannotated splicing events, we manually queried RUM alignments from a subset of our data, including mouse diaphragm, tongue, abdominal aorta, *EDL* and *soleus*. (A) The number of unique novel splicing events detected at different read depths in different mouse muscle tissues is plotted as a bar graph. (B) We manually curated the highest confidence hits (supporting reads > 100) and identified five major classes of novel splicing events. Relative proportions are represented as a pie chart; the most common novel splicing event were novel exons. Panel (C) shows a cartoon diagram of *Myosin light chain kinase 4 (Mylk4)* splicing between canonical exons 2 and 3 (dark grey boxes). RNA-seq data predict the presence of two heretofore unannotated exons, which we provisionally term exons 2.1 and 2.2 (light grey boxes). Black lines represent canonical splicing; blue lines represent potentially novel splicing events. Red arrows indicate the location of RT-PCR primers designed to verify whether these predicted events occur in muscle tissue. (D) RT-PCR gel confirming that the novel splicing events occur in vivo. Input RNA was taken from male mouse *EDL* and *soleus* muscle *collected independently* of the samples used for RNA-seq. Experimental lanes 1 and 2 using primers spanning canonical exons 2 and 3, show a predominant PCR product of ~880bp in both *EDL* and *soleus,* consistent with splicing of both exons 2.1 and 2.2 into the full length transcript. This band was TOPO cloned and conventional Sanger sequencing confirmed the presence of both novel exons. A minor product of ~600bp is also present, which most likely represents the exclusion of one of the two novel exons. Experimental lanes 3 and 4 show a PCR product of ~330bp from both *EDL* and *soleus* using primers that span exons 2.2 to 3, confirming that this splicing event happens in vivo. Experimental lanes 5 and 6 show a PCR product of ~260bp from both *EDL* and *soleus* using primers that span exons 2 to 2.1, confirming that this splicing event happens in vivo. Experimental lanes 7 and 8 show a PCR product spanning canonical exons 7 to 9 of *Mylk4*. These lanes were an internal control for *Mylk4* expression in our samples. Interrogation of the GENCODE database reveals that many of the novel splicing events described above, including those in *Mylk4,* are supported by sequenced cDNAs that have not yet been integrated into gene models.

**Figure S8:**
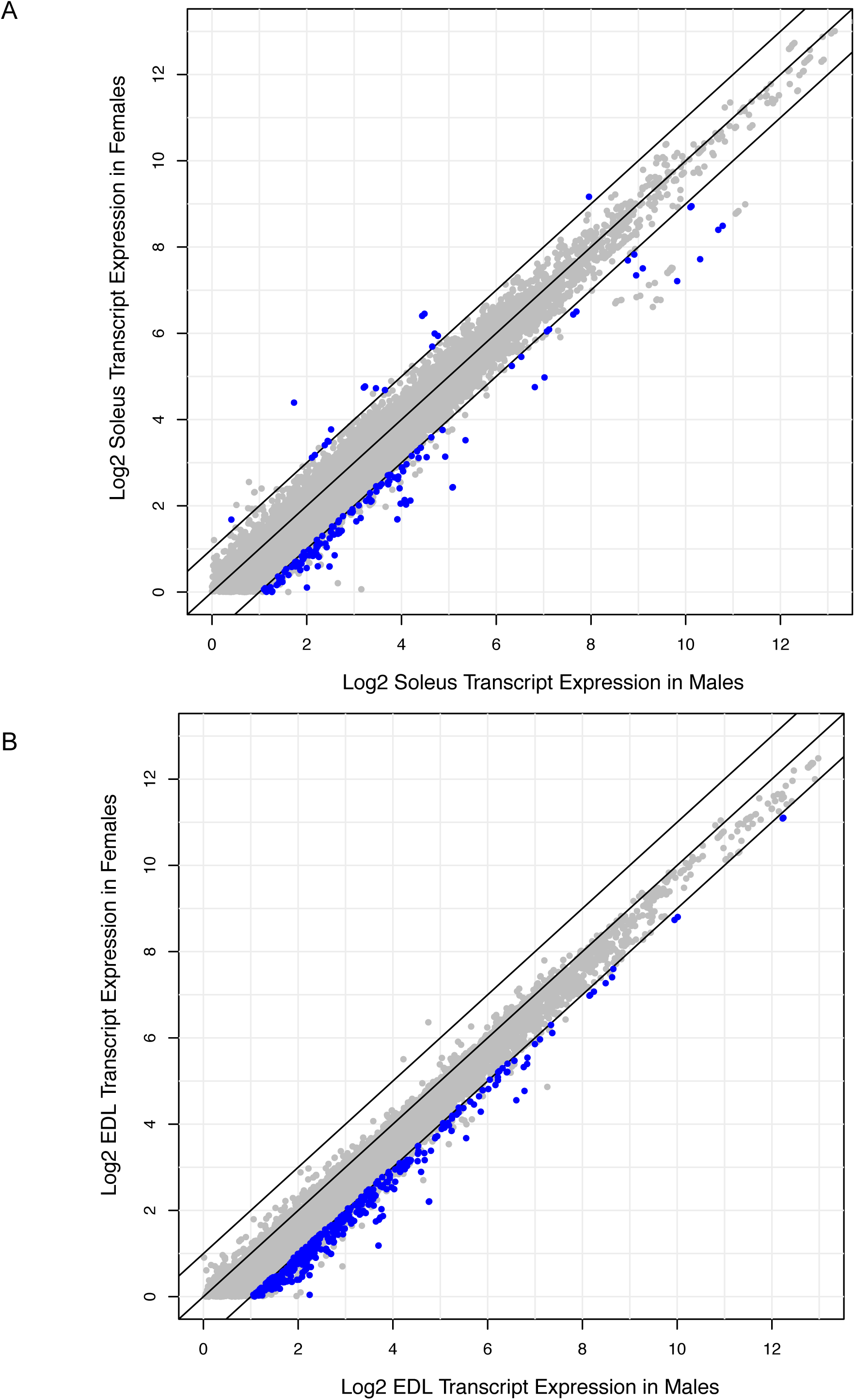
Relatively few genes are differentially expressed between males and females; most of these are up-regulated in males. (A) Scatter-plot showing the log2 expression (FPKM) of every transcript with average expression > 1.0 in rat male *soleus* versus rat female *soleus*. Blue dots represent transcripts with q-values < 0.05 and fold change > 2; grey dots are transcripts that are not statistically significant. Black bars represent fold change cutoffs > 2. (B) Scatter-plot showing the log2 expression (FPKM) of every transcript with average expression > 1.0 in rat male *EDL* versus rat female *EDL*. Blue dots represent transcripts with q-values < 0.05 and fold change > 2; grey dots are transcripts that are not statistically significant. Black bars represent fold change cutoffs > 2.

**Figure S9:**
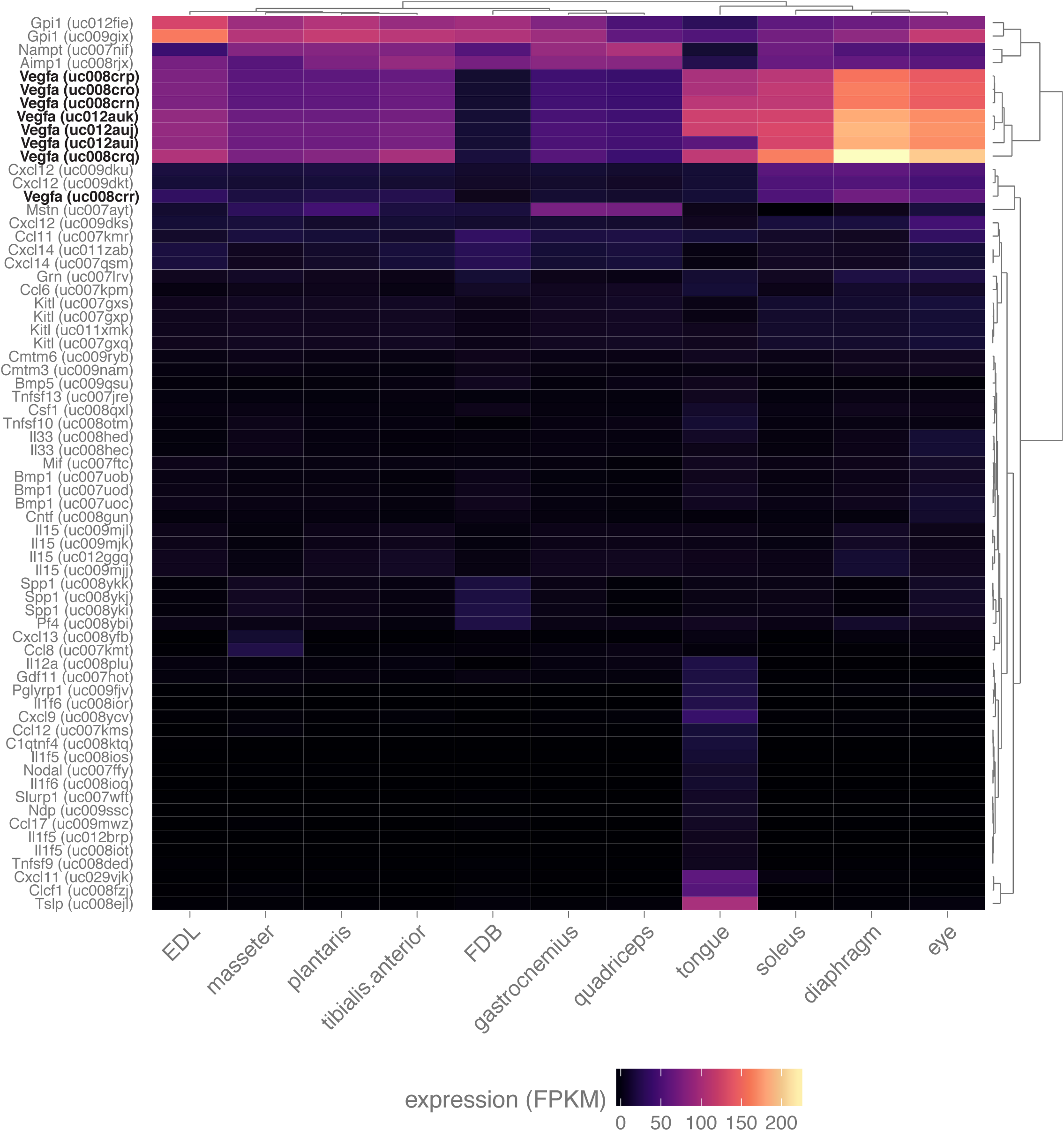
Skeletal muscle expresses numerous myokines. Heat map representation of the expression of all transcripts annotated with the gene ontology term “cytokine activity” and filtered with a q-value threshold < 0.01, minimum expression in at least one tissue > 10 FPKM (N = 67 transcripts). A 10-fold higher expression threshold was chosen for this figure to enrich the gene list for highly-expressed transcripts. Dendrograms represent clustering based on similarity by tissue (top) and transcript (right side). All spliceforms of *Vegfa* (bold, black text) are expressed at notably high levels.

**Figure S10:**
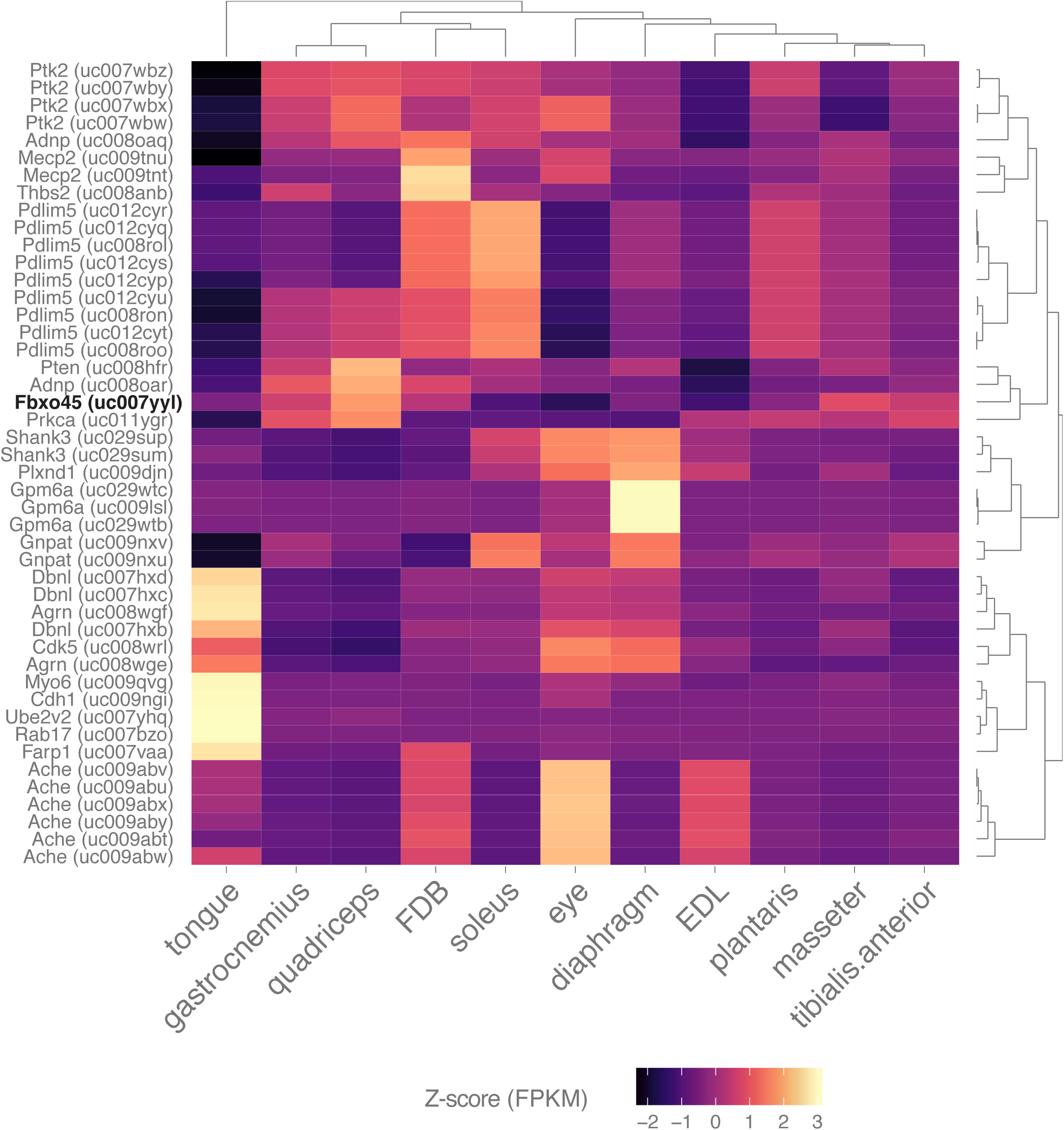
Skeletal muscle expresses many genes involved in synapse assembly. Heat map representation of the expression of all transcripts annotated with the gene ontology term “synapse assembly” and filtered with a q-value threshold < 0.01, minimum expression in at least one tissue > 10 FPKM (N = 46 transcripts). A 10-fold higher expression threshold was chosen for this figure to enrich the gene list for highly-expressed transcripts. Dendrograms represent clustering based on similarity by tissue (top) and transcript (right side). *Fbxo45* (bold, black text) shows differential expression between skeletal muscles and may be involved in neuromuscular junction formation (44).

**Table S1: Details on the total number of reads and aligned reads for every replicate sample.**

**Table S2: Differentially-expressed genes annotated as being involved in skeletal muscle disease.**

**Table S3: Every unique gene that is a candidate myokine.** This Table is distinct from **Figure S9** in that no filtering based on differential expression was performed, and the filter for minimal expression is less stringent (i.e. FPKM > 1 instead of FPKM > 10).

